# Fronto-striatal oscillations predict vocal output in bats

**DOI:** 10.1101/724112

**Authors:** Kristin Weineck, Francisco García-Rosales, Julio C. Hechavarría

## Abstract

The ability to vocalize is ubiquitous in vertebrates, but neural networks leading to vocalization production remain poorly understood. Here we performed simultaneous, large scale, neuronal recordings in the frontal cortex and dorsal striatum (caudate nucleus) during the production of echolocation and non-echolocation calls in bats. This approach allows to assess the general aspects underlying vocalization production in mammals and the unique evolutionary adaptations of bat echolocation. Our findings show that distinct intra-areal brain rhythms in the beta (12-30 Hz) and gamma (30-80 Hz) bands of the local field potential can be used to predict the bats’ vocal output and that phase locking between spikes and field potentials occurs prior vocalization production. Moreover, the fronto-striatal network is differentially coupled in the theta-band during the production of echolocation and non-echolocation calls. Overall, our results present evidence for fronto-striatal network oscillations in motor action prediction in mammals.

## Introduction

Vocalization-based interactions between broadcaster and receiver play an important role in everyday life scenarios and are highly conserved throughout the animal kingdom ^1,2^. Although neural mechanisms involved in auditory processing and perception have been extensively researched ^3–5^, studies addressing subcortico-cortical network activity leading to vocal motor outputs remain sparse.

As bats heavily depend on their ability to vocalize in order to communicate and orient in the environment, they serve as a good animal model for studying the hearing-action cycle. Although bats have been studied for over 50 years ^6^, only a very limited number of experiments were able to obtain electrophysiological recordings from vocalizing bats ^7–9^. So far, most research investigating network activity leading to vocal output focussed on humans ^10,11^. Studies in humans typically use non-invasive measuring techniques that do not allow to link oscillations with spiking activity, especially in subcortical regions.

In this paper, we used the bat species *Carollia perspicillata* to investigate the involvement of fronto-striatal network activity in vocalization production. This bat species belong to the Microchiropterans which are characterized by laryngeal echolocation, similar to human laryngeal-based speech production^12^. *C. perspicillata’s* calls can be broadly split into two types of outputs including echolocation, which typically contain carrier frequencies in the range of 60-90 kHz, and non-echolocation calls such as communication, distress and social calls, whose energy peaks at frequencies below 50 kHz ^13^. Whether differences exist in the neuronal activity patterns leading to the production of echolocation and non-echolocation call remains unknown.

Fronto-striatal networks are a good candidate for studying the mechanisms leading to vocal production. They connect different parts of the frontal lobe with various regions of the striatum which constitutes a major input structure into the basal ganglia ^14,15^. Using tractorgraphic methods, a direct connection between the (dorsolateral) caudate nucleus (CN) with the dorsolateral prefrontal cortex (PFC) forming the “associative circuit” has been identified ^16,17^. Although functions of this circuit in working memory and executive function have been demonstrated ^15^, literature addressing its functional role in self-initiated motor movement, such as vocalization production, is scarce. Morphological alterations in the fronto-striatal path have been observed in diseases accompanied with speech impairment such as Huntington Disease, Parkinson or Asperger Syndrome ^15,18^. The latter could suggest the involvement of frontal and striatal areas in mediating and predicting vocal output. Moreover, studies examining frontal and striatal regions, have identified their putative role in vocalization production in humans ^19–21^and bats ^22^.

In bats, the frontal lobe is a rather unexplored region. Most previous experiments evaluated the auditory responsiveness of the frontal cortex and defined the frontal auditory field (FAF) ^23–25^. It remains controversial whether the FAF is an analogue to the prefrontal areas found in other mammals based on morphology and connectivity ^26,27^. This work will refer to the FAF when discussing the recordings from the bats’ frontal lobe. We hypothesized that the production of echolocation and non-echolocation calls could involve different fronto-striatal network dynamics. This hypothesis was based on the fact that echolocation is used to create an acoustic image of the environment (which depend on listening to echoes of the calls emitted), while non-echolocation vocalizations are uttered to convey information to other individuals. Our findings confirmed this hypothesis and indicate that distinct brain rhythms shape the bats’ vocal output. These rhythms encompass inter-areal coupling in the theta band (4-8 Hz) and specialized intra-areal processing mechanisms in the gamma (30-80 Hz) and beta (12-30 Hz) bands of the local field potentials (LFP). Overall, our results present evidence for fronto-striatal network oscillations in motor action prediction.

## Results

To assess fronto-striatal network activity during vocalization production, 47 extracellular, paired recordings were acquired from the FAF and the caudate nucleus of the dorsal striatum of four male bats. Striatal recordings were performed with linear tetrodes (electrode spacing: 200 μm) while FAF activity was measured with linear 16-channel probes (electrode spacing: 50 μm). The placement of chronically implanted tetrodes in the CN was confirmed histologically for each animal (see example Nissl section in Supplementary Fig. 1). The laminar probe used for FAF measurements was introduced on each recording day. Throughout the manuscript, we will refer to different frequency bands of the local field potentials as theta (4-8 Hz), alpha (8-12 Hz), low beta (12-20 Hz), high beta (20-30 Hz), low gamma (30-50 Hz), and high gamma (50-80 Hz).

### Properties of bat vocalizations

Individual bats were placed in an acoustically and electrically isolated chamber and let to vocalize spontaneously while neural activity in the CN and FAF were simultaneously measured. A total of 39014 spontaneously emitted calls were recorded from implanted, head-fixed animals. Most of the vocalizations produced occurred as trains of syllables produced at short intervals (Fig. 1a, b). Across recordings, the median calling interval amounted to 12 ms ± 54 ms (± inter-quartile range; IQR). For analyzing neural activity related to vocalization production (see below), only utterances surrounded by at least 500 ms pre- and post-time without sounds were chosen. A pool of 628 non-echolocation and 493 echolocation calls remained after vocalization selection (non-echolocation: 628/16204 (3.9 %) and echolocation: 493/22810 (2.2 %)). The main criterion used for classifying sounds into echolocation and non-echolocation was based on their spectro-temporal structure. It is known that *C. perspicillata’s* echolocation calls are downward frequency modulated and peak at high frequencies > 50 kHz (see example spectrogram in Fig. 1c), while non-echolocation calls contain most energy at lower frequencies, generally below 50 kHz (see examples in Fig. 1d, e) ^13^. At the population level, call durations of both types of isolated vocalizations were only slightly different (p= 0.02, Wilcoxon rank-sum test, Fig. 1f, h, the test considered only the temporally isolated calls used for further analysis) with a median around 0.5 ms in both cases but with IQR values of ± 0.8 ms for non-echolocation and ± 1.8 ms for echolocation. As expected, peak frequency differed significantly between the two call types (echolocation: 75.1 kHz ± 32.2 kHz and non-echolocation 12.6 kHz ± 33.2 kHz, Rank-sum Test p < 0.0001, Fig. 1g, i). As peak frequency was used as the main distinctive feature for characterizing the two call classes, the results described in the preceding text constitute a proof-of-principle.

**Figure 1.**
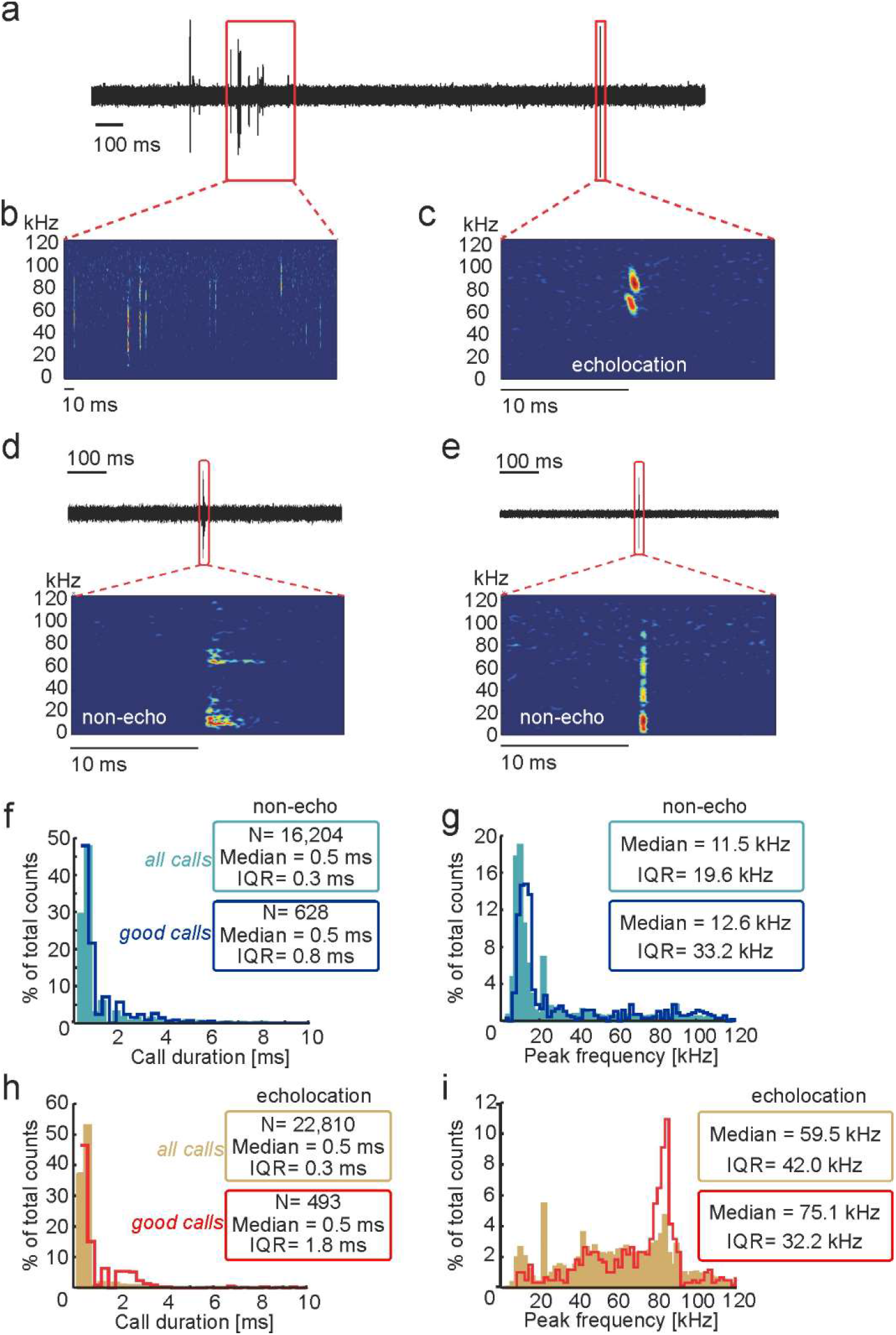
Properties of echolocation and non-echolocation calls produced by bats. **a,** Exemplary acoustic recording including an isolated call and a syllable train. Zoomed-in views have been included in **b** and **c** to show spectrograms of the syllable train and the isolated echolocation call (b and c, respectively). **d** and **e**, show two further examples of isolated vocalizations (non-echolocation calls in this case). **f** and **g** show histograms of call duration and peak frequencies, respectively, for all non-echolocation calls recorded (light blue) and the isolated non-echolocation calls (dark blue; n=628, labeled as “good calls”). The latter were considered for further analysis. In each histogram the median and the interquartile range values (IQR) are given. **h**, **i**, Same as f, g but for echolocation calls. Echolocation and non-echolocation vocalizations differed only slightly in their duration (p-value=0.02, Wilcoxon rank-sum test) but differed markedly in their peak frequencies (p<0.0001).

Besides spontaneous vocalization production, bats were presented with pure tones (10 - 90 kHz in steps of 5 kHz at 60 dB SPL with 10 ms duration) to evaluate auditory responsiveness in the neural populations recorded (see frequency tuning results in Supplementary Fig. 2). The acquired LFPs in both brain regions showed pronounced responses to sounds (see population evoked responses in Supplementary Figure 2a, b and i-k), revealing a preference towards low frequencies around 15-20 kHz (best frequency distributions for both structures studied are shown in Supplementary Fig. 2c, d). Within columns of the FAF, channels located at depths below 400 μm showed the highest auditory responsiveness and neighboring channels had similar frequency tuning properties (see comparison of frequency tuning curves across cortical layers in Supplementary Fig. 2e).

### The spectral structure of LFPs predicts vocal output

LFPs occurring 500 ms before and after call onset were analyzed to gain insights into the involvement of fronto-striatal regions in vocalization production. LFPs were filtered (1-90 Hz) demeaned and z-normalized (see methods). Average LFPs obtained in the CN and FAF are shown in Figure 2 (CN: Fig. 2a, b; FAF: colormaps in Fig. 2c, d; see also supplementary Figure S3 for recordings in one example column). Deflections in the LFP signals following the production of echolocation and non-echolocation calls were evident in both brain areas. These deflections may reflect evoked responses related to the processing of the vocalizations produced. In the FAF, vocalization-evoked responses were strongest in deep layers (i.e channels located at depths >400 μm) matching also the areas of highest responsivity to pure tones (compare colormaps in Fig. 2 c, d with the colormap in Supplementary Figure S2, see also the example column in Supplementary Figure S3). In both brain structures, LFP deflections preceding vocalization production were also observed, although their amplitude was lower than that of vocalization-evoked responses.

**Figure 2.**
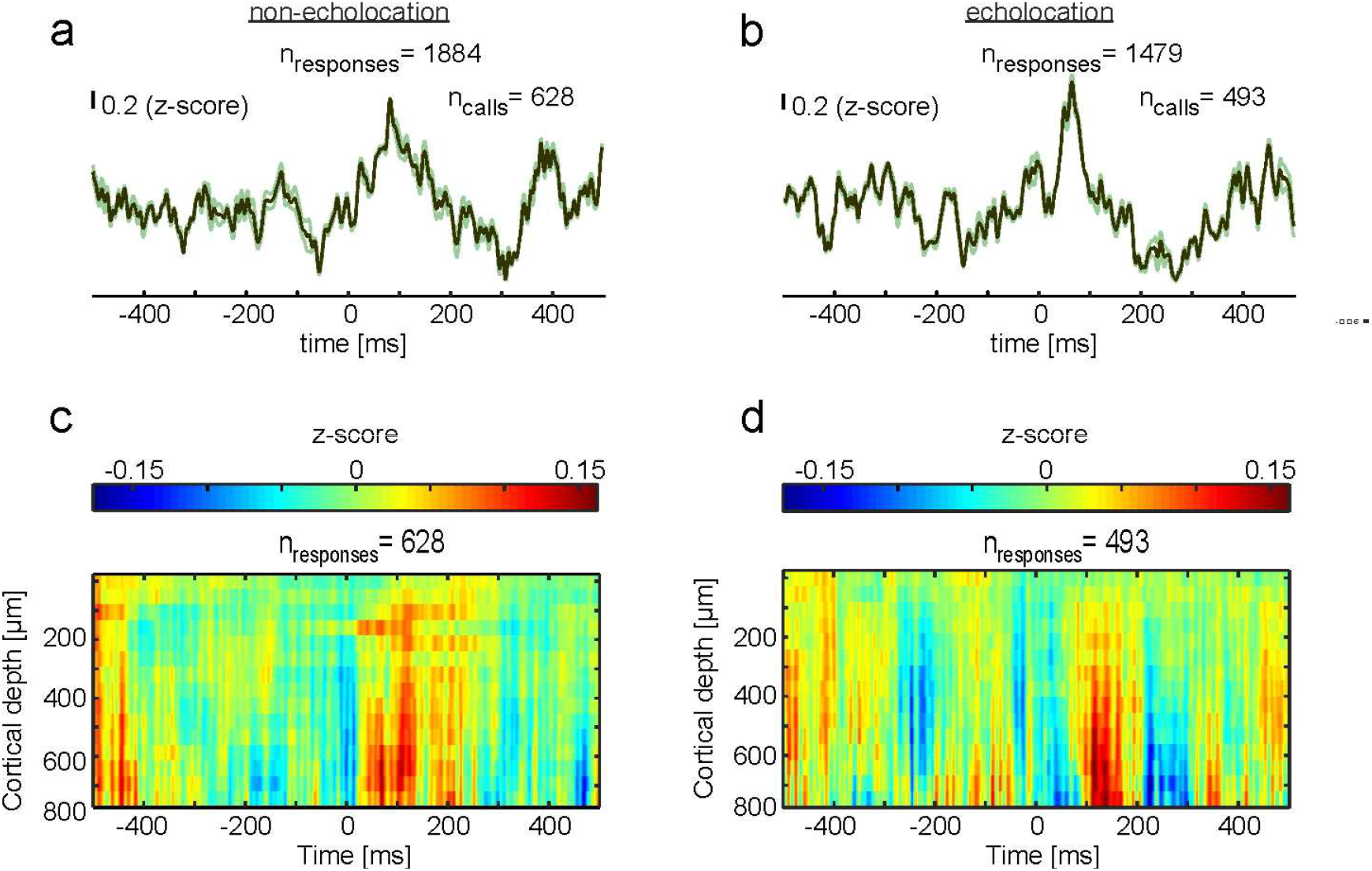
Local field potentials during vocalization production in the CN and FAF. **a,** Mean LFP (± SEM) of all recorded isolated non-echolocation calls (n=628). Signals from all three channels of the striatum were pooled together thus rendering a higher number of responses for the striatum than for the FAF. **b**, Mean LFP (± SEM) obtained during the production of isolated echolocation calls (n=493) in the striatum. **c** and **d**, Colormaps showing the mean of z-scored LFPs in the FAF across cortical depths, 500 ms before and after non-echolocation (c) and echolocation production (d).

Next, we performed spectral analysis of the LFP signals. LFP spectrograms were calculated from bootstrapped signals based on 10,000 randomization trials for each vocalization type (see methods). This approach allows to assess spectral components that are consistently time-locked across vocalization trials. The striatal spectrograms followed the typical power rule by which high power occurred in the low LFP frequencies and power decreased as LFP frequency increased (Fig. 3a, b). When comparing both conditions (echo vs. non-echo) with each other, time and frequency dependent variations could be detected. These differences became obvious when comparing both power spectrograms using the Cliff’s delta (*d*) metric (Fig. 3c). This metric computes the effect size of group comparisons and ranges from −1 to 1 with identical groups rendering values around zero ^29^. Cliff’s Delta matrices revealed higher power in the gamma range of the LFP (especially frequencies >70 Hz) before non-echolocation production than prior echolocation (blue areas in Fig. 3c). Differences in the gamma range prior vocalization had a medium-size effect (grey contour lines in Fig. 3c) following values proposed in previous studies ^29^. In contrast, power in the beta range (12-30 Hz) was found to be more pronounced before and during echolocation than during non-echolocation. Both observed effects suggest that different striatal LFP frequencies play different roles in vocalization production (beta higher for echolocation and gamma for non-echolocation). To portray the power of individual examples, representative single trials of LFP signals in the frequency ranges displaying the highest vocalization-dependent differences are shown in Supplementary Figure 4a-d (striatum) and e-l (FAF).

**Figure 3.**
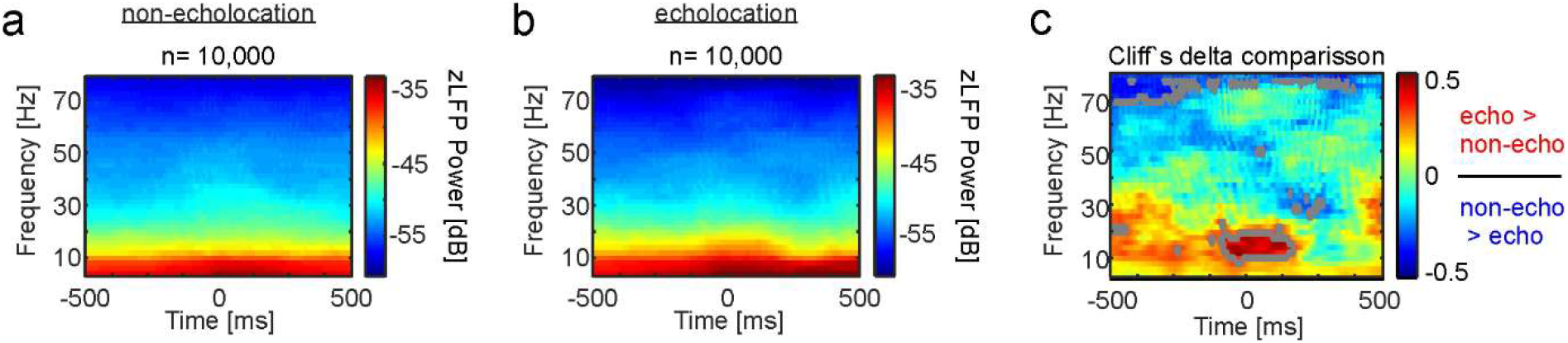
Spectral differences in neural activity obtained in the CN during echolocation and non-echolocation production. **a-b,** Power spectrogram in the CN during non-echolocation (a) and echolocation (b). Mean values of 10,000 randomization trials (see methods) are displayed in each case. **c**, Colormap representing the Cliff’s Delta values of echolocation vs. non-echolocation comparisons at each time point and frequency. Grey outlined regions mark areas with a medium effect size (Cliff Delta > 0.33 ^29^). Red colours indicate more power in the LFPs during echolocation than non-echolocation, whereas blue regions indicate the opposite trend.

Similar to the CN, spectrograms of FAF neural signals related to non-echolocation (Fig. 4a-d) and echolocation (Fig. 4e-h) calls followed a power rule. Large differences could be detected when comparing the neural spectrograms obtained during echolocation and non-echolocation (Fig. 4i-l). The largest differences were found in the low- and high-gamma range (30-50 and 50-80 Hz, respectively) with the power being higher before and during echolocation than during non-echolocation production, especially at FAF depths below 200 μm. The latter is illustrated in Figure 4i-l for four example recording channels located at different depths and in Figure 4o for all FAF depths studied. Other large spectral differences were found in the theta-alpha range (4-12 Hz) both before and after vocalization production with a time- and depth-dependent pattern (see red and blue regions in example channels in Fig. 4i-l and across-depths data Fig. 4m). Differences in the beta band (12-30 Hz) were pronounced mostly before sound production and occurred at different time points before call onset across cortical depths (Fig. 4n). Overall, these results suggest that different neural frequency channels in the FAF and CN contribute differently to the bats vocal output.

**Figure 4.**
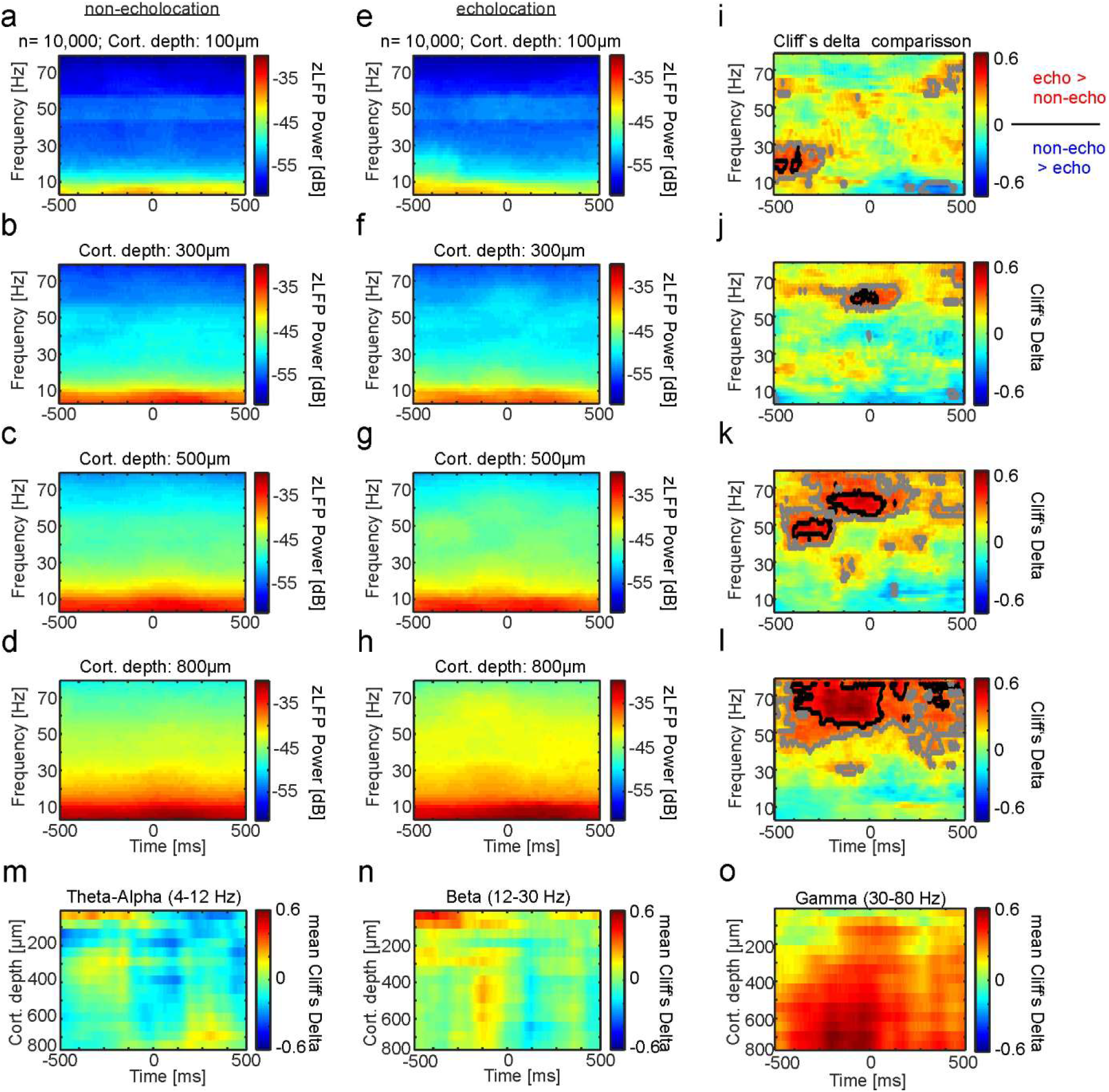
Time-frequency differences in power distributions across FAF channels depending on the vocalization type. **a-d,** LFP spectrograms of four illustrative channels of the FAF for the non-echolocation condition (n= 10,000 randomization trials, see methods). **e** - **h** Spectrograms obtained in the same four example channels during echolocation production. **i-l**, Colormaps of Cliff’s Delta values obtained when comparing the time-frequency dynamics in the echolocation and non-echolocation conditions in the four example channels. Black highlighted regions indicate large effect size (>0.47). Grey indicates medium effect size (>0.33)^29^. **m-o** Mean Cliff’s Delta values across FAF depths. Mean values were obtained for all the frequencies that composed the theta (4-8 Hz), beta (12-30 Hz), and gamma bands (30-80 Hz), represented in panels m, n and o, respectively.

We used binary support vector machine (SVM) classifiers to assess whether models could be constructed to ‘predict’ the bats’ vocal output based solely on the power distribution of LFPs before call production. SVM classifiers were trained (only once) with 5,000 randomly chosen power distributions across time per vocalization type and frequency band (for each frequency band the average power at each time point was calculated). As mentioned, only spectral power occurring before call onset was considered for training and predicting vocal output. The remaining 10, 000 power distributions (5,000 per call type) were used to compute the percentage of correct hits by the models (Fig. 5a, b). In the CN, LFPs especially the low beta (12-20 Hz) and high-gamma (50-80 Hz) bands, provided the best predictions about the type of upcoming vocal outputs (~65% correct hits in both cases; Fig. 5a). Note that these frequency bands showed the highest differences in power when comparing both vocalization conditions (cf. Fig. 3c). Overall, the FAF provided higher prediction accuracy than the CN (Fig. 5b). Here, the gamma band (in particular the high gamma band (50-80 Hz)) displayed a large accuracy in predicting the type of vocal output reaching values ~80 % accuracy at depths >500 μm. Gamma signals in the FAF also produced the lowest model cross-validation errors (see Supplementary Figure S5a, b). Note that in both the CN and FAF, training the SVM classifiers with false information, created by randomization of the labels in training signals, led to a drop in prediction capability with true detection rates around chance level (i.e. 50 % see Supplementary Figure 5c, d).

**Figure 5.**
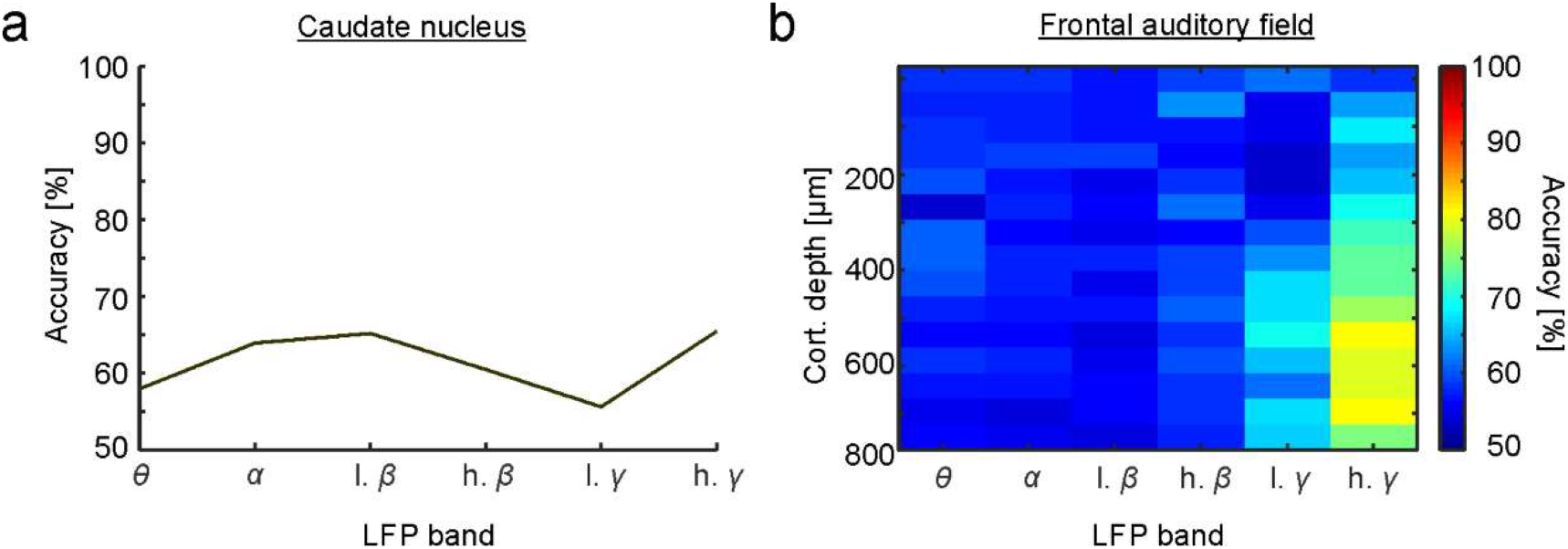
LFP signals leading to vocalization can be used to predict vocal output. **a,** Prediction accuracy was calculated for models built using a binary support vector machine classifier (see methods), trained with LFP information (filtered by frequency band) occurring before vocalization production in the echolocation and non-echolocation conditions. Models were trained with half of the data (n=5,000 randomization trials in each vocalization condition). The other data half was used for calculating the models’ prediction accuracy. **a**, prediction accuracy for the CN. **b**, prediction accuracy for the FAF. Both brain structures showed the highest prediction accuracy in the high gamma range and the FAF rendered better predictions than the CN.

### Fronto-striatal coupling occurs in low frequency bands of the LFP

To investigate the functional coupling between the FAF and CN during vocalization production, the neural “coherency” was calculated. Coherency refers to the trial-averaged cross-spectral density of two signals measured simultaneously, taking into account the phase synchrony of the signals ^30^. Here, the magnitude of coherency (defined as “coherence”) was calculated between neural signals recorded at different depths of the FAF and the CN (Fig. 6). The preferred frequencies for coherence between both structures were located in the low spectral range (under 12 Hz, mostly in theta (4-8 Hz), see below) for both types of vocalizations. There was a striking difference in the temporal pattern of coherence observed in the two vocalization conditions. For non-echolocation calls, the highest fronto-striatal coherence was found before calls were uttered (Fig. 6a-d). However, when echolocation-calls were produced, coherence shifted to the time points after call emission (Fig. g-j). The different temporal coherence patterns in the two vocalization conditions were also clear in average coherence plots that display the mean theta and alpha coherence across all FAF depths studied (Figure 6e, f and k, l). Note that regardless of the vocalization type produced, FAF depths below 600 μm rendered the lowest coherence values even though they displayed the strongest LFP deflections (compare results in Fig. 6e and k with Fig. 2c and d). Also note that the gamma band of the LFP (>30 Hz) was not involved in inter-areal coherence, even though this band did show differences in within-structure analysis of LFP signals during echolocation and non-echolocation production (see results presented in Figs. 3 and 4). Taken together ours results indicate temporally defined functional coupling of fronto-striatal circuits depending on the type of the vocal output produced by bats.

**Figure 6.**
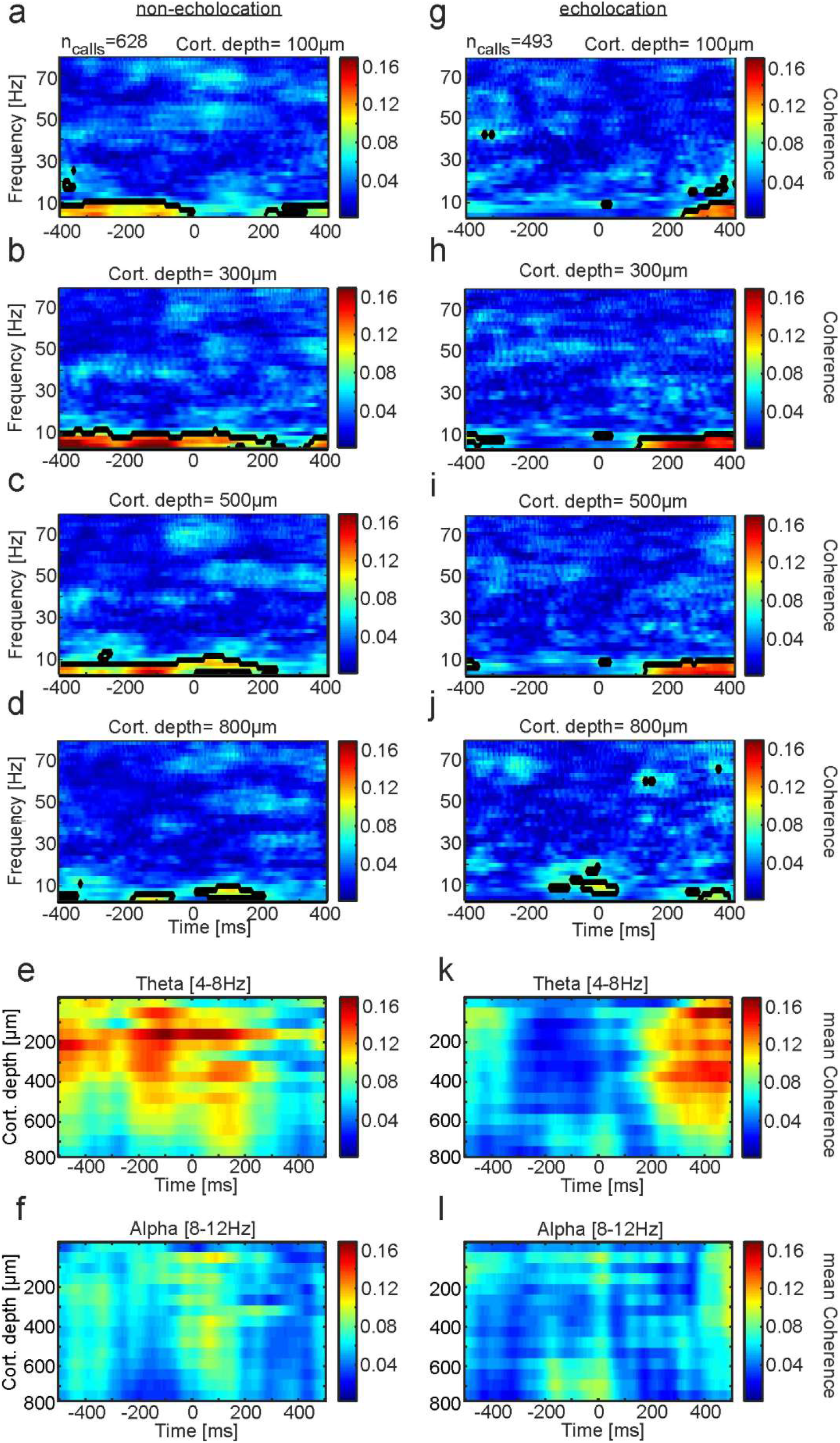
Functional coupling between the FAF and the CN during vocalization production. Time-frequency resolved coherence between the striatum and four exemplary channels of the FAF at: **a**, 100 μm; **b**, 300μm; **c**, 500μm; and **d**, 800μm depths during non-echolocation (n=628 trials). Black coloured regions refer to the 95^th^ percentile of all computed coherence values during vocalization production. **e**, Time resolved coherence strength between both structures across cortical depths in theta (4-8 Hz), and alpha (8-12 Hz; panel **f**) during the production of non-echolocation calls. Mean coherence values across frequencies included in each range were calculated. **g-j** Coherograms in four example channels during echolocation call production (n=493 trials). **k**, **l**, Same as e, f but for the echolocation case. During echolocation production pronounced coherence in theta in the top-to-middle layers was found 200 ms after call onset. This temporal pattern differs from that observed during the production of non-echolocation calls.

### Frequency dependent spike-LFP locking prior vocalization production

We also studied the spiking pattern of striatal and FAF neurons and the relation between spiking and LFP phase within and between structures. Spiking activity was gathered from spike-sorted single units (see methods). Peri-stimulus time histograms (PSTHs) computed for the CN did not show clear evidence for evoked responses following vocalization production in either vocalization condition (Fig. 7a, b). In the FAF, spiking was strongest in superficial and deep layers and vocalization-triggered spiking was apparent at depths below 600 μm in both vocalization conditions (Fig. 7c, d).

**Figure 7.**
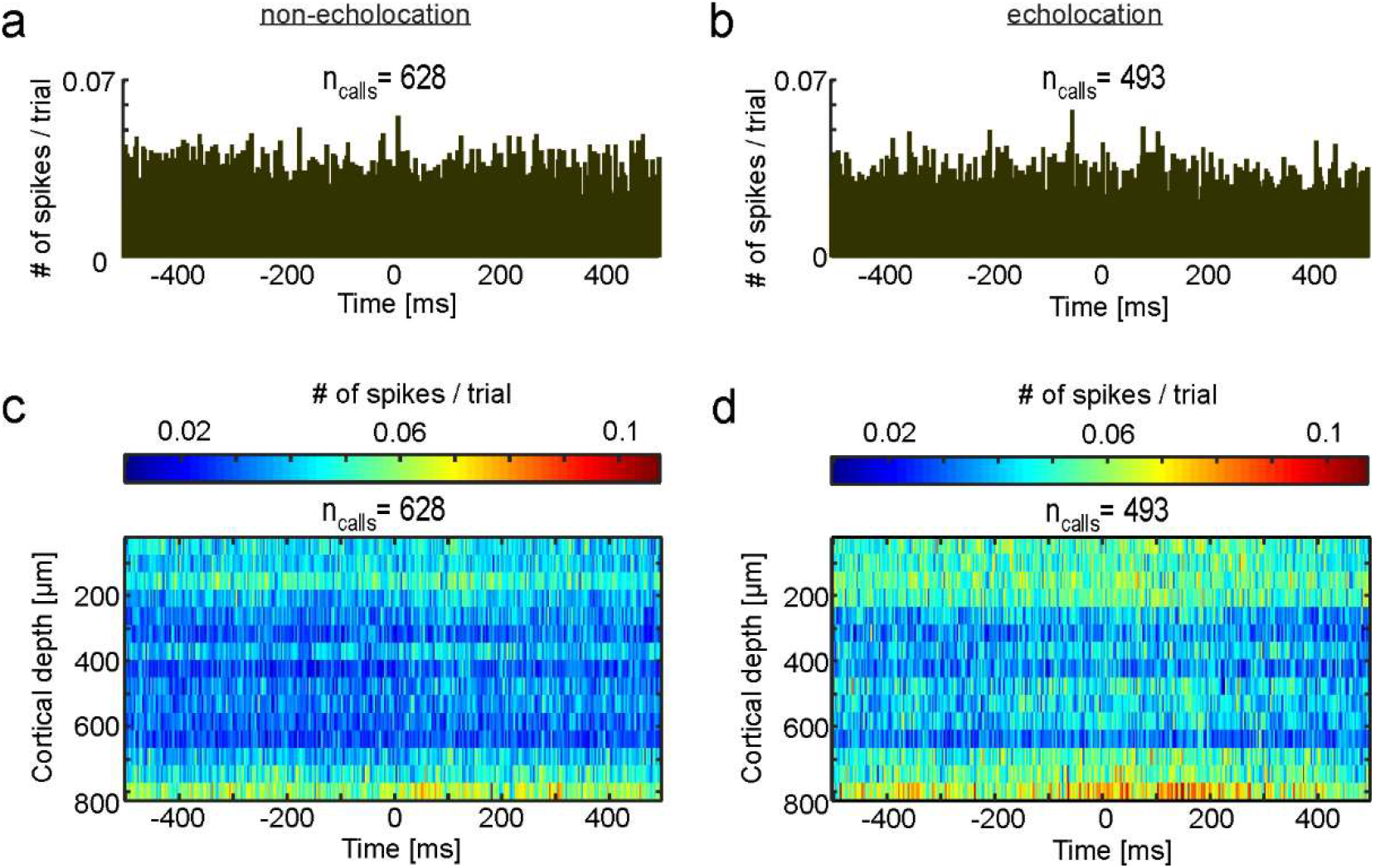
Spiking activity in the CN and FAF during vocalization production. Spiking probability (computed as numbers of spikes per trial per bin, binsize=3 ms) in the CN 500 ms before and after non-echolocation (**a**, n=628 trials) and echolocation (**b**, n=493) production. **c-d**, Spiking across all 15 channels recorded in the FAF during non-echolocation (**c**) and echolocation (**d**). In the FAF, during both types of vocalizations, distinct spiking activity could be identified in deep layers.

Next, the locking between spikes and the phase of LFPs occurring before vocalization production was studied. Phase-locking values were calculated by linking spike-times to the instantaneous phase of each LFP frequency band (see example phase locking calculations in Supplementary Figure S6). The circular distributions of LFP phases at which spiking occurred for each frequency band were compared with random-phase distributions obtained by extracting LFP phases at time points not related to spiking. To obtain robust circular spike-phase and random-phase distributions, circular distributions were calculated 10,000 times, with 100 randomly chosen spike-phase and random-phase values included in each randomization trial (see methods). Differences in vector strength (dVS) between spike-phase and random-phase distributions were calculated to estimate the strength of spike-phase locking (see circular distributions and vector strengths (VS) in supplementary Figure S7). Significance was assessed by comparing VS values obtained across randomization trials for the spike-phase and random-phase conditions (Bonferroni corrected Wilcoxon rank-sum test p<0.001, see methods).

In the CN, significant differences between spike-phase and random-phase distributions were found in the theta band during non-echolocation, and in the alpha and high-gamma bands during echolocation (Fig. 8a, b). When comparing VS distributions obtained in both vocalization conditions (not with the surrogate data) in the striatum, significant differences were only found in the high beta range (Fig. 8c). The FAF showed statistically significant spike-phase locking in several LFP frequency bands and cortical depths (Fig. 8 d, e). In particular, spike-phase locking in the low- and high-gamma LFP bands was pronounced across layers and consistent differences when comparing between vocalization types appeared in the low-gamma range at FAF depths >600 μm (Fig. 8f). Besides the gamma spike-phase locking observed, in the non-echolocation condition, there was consistent spike-phase locking in the theta band at depths spanning from 250-400 μm (Fig. 8d), although this effect was not significant when comparing between vocalization types (Fig. 8f). Note that we refer to “consistent” spike phase-locking differences whenever statistical significance occurred in more than two contiguous FAF channels. Also note that effect size calculations (e.g Cliff’s delta) complementing rank-sum testing rendered in all cases values below 0.2, thus indicating high data variability (see effect size plots in Supplementary Figure 8).

**Figure 8.**
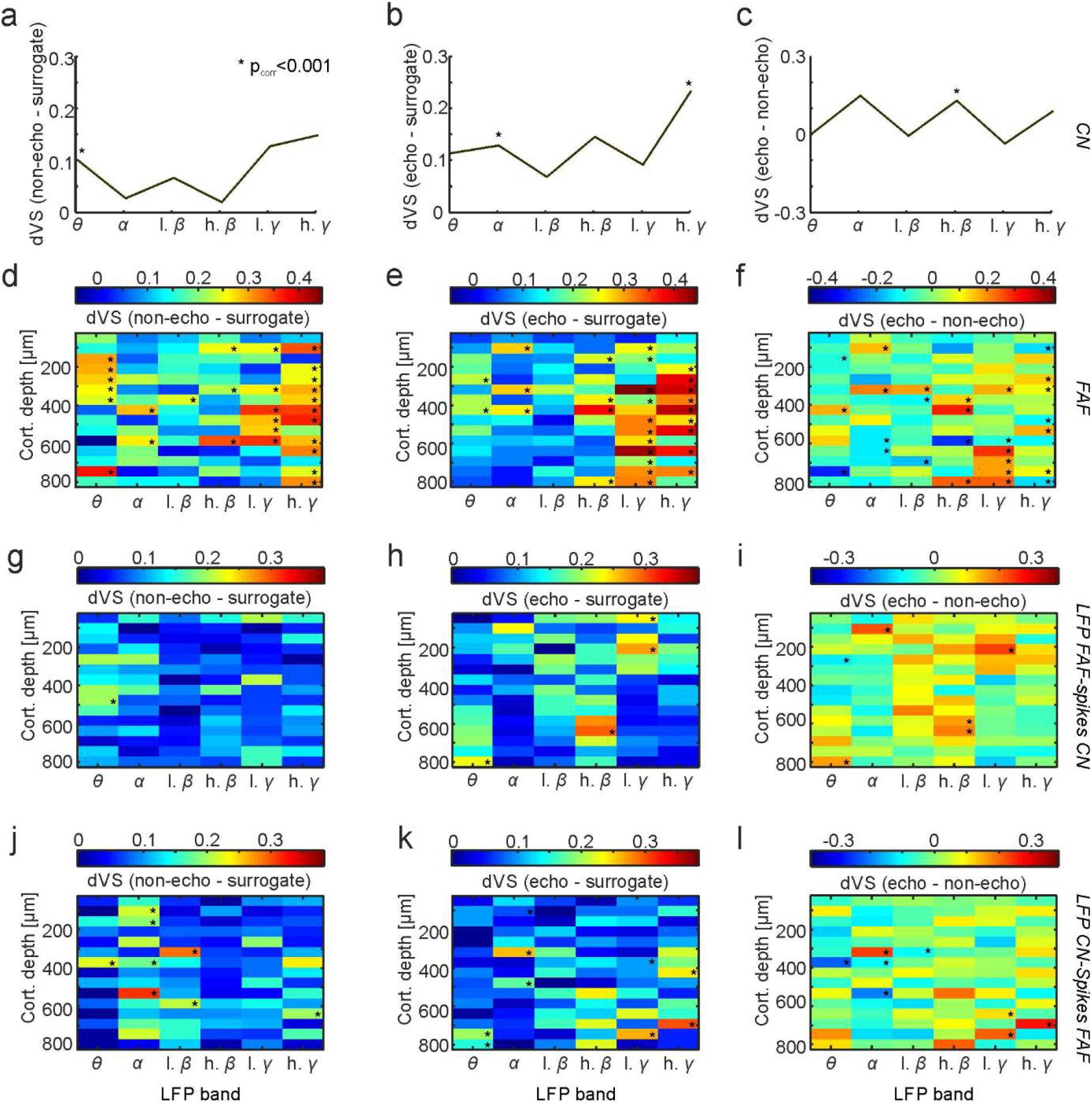
Spike phase locking across vocalization conditions. **a** Differences in vector strength (dVS) obtained before vocalization onset in the non-echolocation - surrogate condition in the caudate nucleus; **b**, echolocation - surrogate; and **c**, echolocation - non-echolocation. **d-f**, dVS values computed for different channels of the FAF in the three conditions mentioned above. **g-i**, dVS values after linking the phase of LFPs in the FAF to spikes recorded in the CN. **j-l**, dVS values obtained when linking the phase of LFPs in the CN to FAF spikes. Statistical differences were tested by comparing vector strength distributions (Wilcoxon rank-sum tests with Bonferroni correction, * p<0.001, see methods).

To assess the relationship between spikes and LFPs within the fronto-striatal network, phase locking values considering the simultaneously recorded spikes in the CN and the phase of FAF field potentials were calculated (Fig. 8g-i). This approach allows to assess whether inter-areal coupling occurs between spikes and LFPs. Overall, the phase of FAF LFPs appeared to be poorly related to striatal spiking with significance occurring in only few, sparse combinations of LFP-band and FAF depths in all three conditions tested (i.e. non-echolocation vs. surrogate, echolocation vs. surrogate, echolocation vs. non-echolocation, Fig. 8g, h, i, respectively). The same was true when calculating spike phase-locking values by considering FAF spikes and simultaneously recorded striatal LFPs (Fig. 8j, k, l). Based on ours results, we can conclude that inter-areal spike phase locking patterns in the fronto-striatal circuit are less pronounced than the patterns calculated considering intra-areal signals.

## Discussion

Previous work has shown alterations in the fronto-striatal network in diseases with impaired speech production ^15,18^. However, electrophysiological mechanisms by which fronto-striatal activity could participate in vocalization production in healthy mammals remain elusive. In this article, we show that neural oscillations in fronto-striatal circuits are distinctly linked to the type of vocalizations produced by bats. The main findings reported in this paper include; (i) a unique intra-areal pattern of LFP frequency representation during vocalization production (most prominent in beta and gamma LFP ranges (12-30 and 30-80 Hz, respectively) which can be used to predict ensuing vocal actions, (ii) functional coupling between the CN and FAF in low frequencies (theta, 4-8 Hz) with temporally distinct characteristics depending on the vocal output, and (iii) the occurrence of spike-LFP phase locking, especially in frontal areas in the gamma LFP band, prior vocalization. These results suggest a functional involvement of the fronto-striatal network in neural processing of motor action selection and implementation, with the capacity to discriminate between, and predict, different motor outputs. Moreover, the FAF and the dorsal striatum appear to contribute individually on an LFP and single unit basis to vocalization production, but appear to be coupled in distinct frequencies and time points in relation to the motor action. A visual abstract of the results can be found in Supplementary Figure 9.

Our hypothesis that echolocation and non-echolocation production involve different fronto-striatal network dynamics was corroborated. The main differences in producing and processing different types of vocal outputs encompass neural oscillations in the theta, beta and gamma band. To our knowledge, the present study is the first to implement simultaneous electrophysiological recordings from two brain structures in vocalizing bats. So far, only few studies accomplished the combination of electrophysiology with vocalization production in bats ^7–9^. Other studies in freely moving bats have been performed before, but using megachiropterans (“non-laryngeal echolocators”) as animal model ^31–33^.

According to our data, the beta band of the LFP is differentially involved in echolocation and non-echolocation production. Beta power is highest during echolocation production in the CN and in superficial layers of the FAF (see Figs. 3 and 4). In general, beta oscillations are suggested to hold functions in perception, memory and sensory processing ^34–36^. Furthermore, beta has been linked to motor actions in the motor cortex ^37^and other striatal networks involving structures such as the putamen ^38^. Our results, together with those from previous studies, support the view of beta oscillations throughout the dorsal striatum having a crucial role in motor action representation.

Further spectral differences accompanying the production of distinct vocalization types occurred in the theta-alpha range across layers of the FAF. The production of non-echolocation and echolocation calls rendered power in this range after vocalization when analysing LFPs (cf. Fig.4a with Fig. 4b, across-depths data in Fig. 4m) and inter-areal coherence (Fig. 6). Power differences after call onset also occurred in the beta and gamma range (Fig. 4n, o). Note that the time period after vocal production has to be examined cautiously as the calls produced could differ in their acoustic attributes (i.e. frequency composition, duration, among others), which could lead to differences in call-evoked neural responses. In different sensory cortices, low frequencies such as theta and alpha are known to modulate sensory processing and neuronal excitability, facilitate predictive coding and enable sensory selection ^39^. Unlike sensory cortices, low frequency oscillations in frontal areas are less understood in terms of sensory processing.

Our data suggests that low frequency rhythms in frontal areas (especially theta, see Fig. 6) mediate inter-areal communication between superficial layers and the dorsal striatum during vocalization production. This result falls in line with a putative involvement of low frequency oscillations in long range synchrony ^40,41^. The strong theta-band coherence between FAF and CN reported in this study resembles coherence patterns between motor cortex and the dorsal striatum during attentive wakefulness ^42^. The FAF constitutes a non-classical sensory area and its laminar structure (i.e. location of inputs and outputs, such as layer 4 and 5 in sensory cortices ^43^) needs further anatomical exploration. According to our data, FAF layers could hold a crucial role in oscillatory communication between frontal and striatal regions during self-initiated motor movement. When assessing the coupling between fronto-striatal regions, the timing of inter-areal coherence seems to play an important role when planning and producing different types of vocalizations (see Fig. 6). Whereas the highest level of coherence was found before and centered around non-echolocation production, echolocation calls induce coherence at least 200 ms after the calls’ onset. One possible explanation for the latter could be that echolocation calls require a more thorough sensory processing after vocal production (i.e. for echo evaluation) than non-echolocation calls. Such post-processing of echolocation calls could be relevant for successful orientation and a coherent representation of the environment in bats.

Another large vocalization-type dependent effect was detected in the gamma band. Before echolocation, high power in this frequency band was observed in deep layers of the FAF, whereas before non-echolocation high gamma power was found in the CN. As the power maxima in gamma was reversed across vocalization conditions in both brain structures, it could be suggested that each component of the fronto-striatal path relies on a differential power distribution of high frequencies in order to produce the same vocal output. This could be supported by the fact that in both brain structures power in the gamma band was the best predictor of vocal output (Fig. 5). Gamma LFPs also appeared to be related to spiking activity. The time period before both echolocation and non-echolocation call production displayed significant phase-locking values in the gamma range across FAF layers (Fig. 8d, f). The latter suggests a generalized role of gamma-phase coupling preceding vocalization production. Spike-phase locking in the gamma range has been demonstrated previously, linked to vocalization in the sensorimotor nucleus of zebra finches ^44^.

Classical functions of gamma rhythms across species are linked to selective attention, cortical computation and working memory ^45^. In bats, changes in gamma power have been linked to the processing of auditory stimulation in the bat auditory cortex and to social interaction in frontal areas ^31,46^. Moreover, an increase in gamma power was found in the superior colliculus after the production of clusters of echolocation calls in freely flying bats ^7^. The latter could relate to the detected rise of gamma power before echolocation in comparison to non-echolocation in the FAF (this study), and could indicate the putative importance of gamma rhythms during navigation, whether the animals are freely flying (as in previous studies) or exploring their environment using their biosonar from a fixed location (this study). Note that gamma oscillations were not involved in long range fronto-striatal communication (see Fig. 6). This finding supports the current view of gamma rhythms as important mostly for local neural computations ^38,40^.

To our knowledge, changes in gamma power linked to a specific motor action have not been described before for the CN. The ventral striatum is known to display a prominent pattern of gamma power during reward ingestion or decision making ^47^ but the oscillatory properties of the nuclei that form the dorsal striatum (such as the CN) are less studied. Our results show that not only the FAF, but also the CN is able to represent future vocal outcomes based on gamma power. Altogether our findings indicate that neural oscillations in the gamma and beta bands in fronto-striatal brain regions represent ensuing vocal actions in bats, while oscillations in the theta-alpha range represent the differential sensory processing of the type of call uttered and play a role in long-range inter-areal coupling. Our data suggests that neural oscillations are an important component of canonical circuits underlying vocalization production in mammals as well as in evolutionary adaptations supporting bat echolocation.

## Methods

### Animal model

For neurophysiological recordings, 4 adult Seba’s short-tailed bats (*Carollia perspicillata*) were used. The animals originated from a breeding colony at the Institute for Cell Biology and Neuroscience, Goethe University, Frankfurt am Main (Germany), and were treated in accordance with the current guidelines and regulations for animal experimentation and the Declaration of Helsinki. All experiments were approved by the Regierungspräsidium Darmstadt (permit number: FU1226). For experimentation, bats were housed individually.

### Surgical procedure

The surgery encompassed the implantation of a chronic tetrode mounted on a microdrive in the striatum, a craniotomy above the FAF for the insertion of a linear silicon probe, and the attachment of a custom-made metal rod. The latter facilitated stable recording conditions by preventing head movements. The implantation protocol was modified from the procedure previously used ^13^. First, after monitoring the health status, bats were anesthetized subcutaneously with a mixture of ketamine (10 mg/kg Ketavet, Pfizer, Germany) and xylazine (38 mg/kg Rompun, Bayer, Germany) and topically with local anaesthesia (Ropivacaine 1%, AstraZeneca GmbH). After achieving stable anaesthesia conditions, animals were placed on a heating blanket (Harvard, Homoeothermic blanket control unit, MA, USA) at 28°C. Afterwards, the fur on top of the head was excised, the skull was exposed via a longitudinal midline incision and the skin, connective tissue, muscle and debris were removed. Using macroscopically visible landmarks (e. g. the pseudocentral sulcus and blood vessels), the skull was evenly aligned. With a scalpel blade, a first craniotomy (~ 2mm diameter) was made between the sulcus anterior and pseudocentral sulcus for the chronic implantation of a tetrode (Q1-4-5mm-200-177-HQ4_21mm, NeuroNexus, Ann Arbor, MI, USA, see supplementary Figure 1) mounted on a moveable microdrive (dDrive-m, NeuroNexus, Ann Arbor, MI, USA) to ensure mobility of the electrodes. To prevent the electrodes from bending, the tetrode was introduced into the tissue (partial implant with 2.0 mm depth) with an angle of 17° perpendicular to the brain surface under the microscope. Subsequently, the microdrive was fixed to the scalp with a two component UV-acrylic glue (Kulzer GmbH, Hanau, Germany) and dental cement (Paladur, Kulzer GmbH, Hanau, Germany) and was placed via a screw (1 full counter-clockwise turn = 150μm) at the target position (in total: >2.1 mm depth). For protection and shielding, a plastic cap covering the implant was glued using UV-acrylic. The connector was permanently attached to the cap. For stability purposes, a custom-made metal rod (2cm length, 0.1 cm diameter) was fixed to the surface of the bat’s skull. The metal post was glued using UV acrylic and dental cement to the bone and the plastic cap posterior to the tetrode (see sketch in Supplementary Fig. 1A). All efforts were made to reduce the weight of the implant.

Before starting the experiments, a second craniotomy (~2-3 mm diameter) rostral to the tetrode between the sulcus anterior and longitudinal fissure above the FAF was implemented using a scalpel blade ^23^. To record extracellular action potentials and LFPs in the FAF, an acute A16 laminar probe (NeuroNexus, Ann Arbor, MI, USA, Fig. 1B) was introduced into the brain on each recording day. After surgery, the animals had at least 48 hours of recovery before starting electrophysiological recordings.

### Neurophysiological recording in the vocalizing animal

All experiments were performed chronically for a maximum of 2 weeks after surgery. Whenever the wounds were handled, local anesthesia (Ropivacaine 1%, AstraZeneca GmbH) was administered topically. Before starting the electrophysiological recordings, the bat was placed in a custom-made holder with an attached heating blanket (see previous section) in a Faraday chamber. Subsequently, the tetrode was connected via an adaptor (Adpt. CQ4-Omnetics16, NeuroNexus, Ann Arbor, MI, USA) to a micro amplifier (MPA 16, Multichannel Systems MCS GmbH, Reutlingen, Germany). For detecting neural activity in the FAF, the laminar probe was lowered through the craniotomy under the cortical surface using a micro manipulator (piezo manipulator PM101, Science Products GmbH, Hofheim, Germany) with a speed of 50 μm/s. The linear probe spanned cortical depths of 50-800 μm below the brain’s surface, with channels evenly distributed in 50 μm steps. One silver wire was placed above the dura mater through a third small craniotomy and served as common ground electrode for both the tetrode and the laminar probe. The reference of each electrode array was short-circuited with the respective top recording channel (the electrode closest to the brain surface) to obtain local signals and prevent movement artefacts. Neuronal signals from the striatum and FAF were preamplified and connected via flexible cables to a portable multichannel recording system with integrated AD converter (Multi Channel Systems MCS GmbH, model ME32 System, Germany). The recording was digitized at a sampling frequency of 20 kHz (16-bit precision). For monitoring, visualizing and storing the data, MC_Rack_Software Version 4.6.2 (Multi Channel Systems MCS GmbH, Reutlingen, Germany) was used.

For the acquisition of vocal outputs, a microphone (CMPA microphone, Avisoft Bioacustics, Glienicke, Germany) was placed 10cm in front of the animal. Acoustic recordings were conducted with a sampling rate of 250 kHz. Vocalizations were amplified (gain = 0.5, Avisoft UltraSoundGate 116Hm mobile recording interface system) and stored in a PC using the Avisoft Recorder Software (Avisoft Bioacoustics, Glienicke, Germany) with 16 bit precision. Offline analysis was conducted to separate vocalizations into echolocation and non-echolocation calls based on their spectro-temporal structure.

In order to synchronize the recording of the vocalization signals and the neurophysiological signals, Matlab-generated triggers (i.e. a sound for acoustic recordings and a TTL pulse for the neural acquisition system) were used to align both recordings. Each recording comprised 3 × 10 min vocalization experiments during which bats were let to vocalize at their own volition, with a short break to stimulate vocal production by opening and closing the recording chamber.

### Acoustic stimulation

To estimate the responsiveness of the areas studied to acoustic stimuli, a frequency tuning paradigm was used. Frequency tuning was controlled via a custom-written Matlab software (Math Works, Natick, MA, USA). A stimulation speaker (NeoCD 1.0 Ribbon Tweeter; Fuontek Electronics, China) was placed 12 cm in front of the animal and pure tones were presented ranging from 10 – 90 kHz in 5 kHz steps (randomized order, repetitions of each pure tone= 30 times) for a duration of 10ms (0.5ms rise/fall time) at 60 dB SPL. Following digital-to-analogue conversion using a soundcard (RME Fireface 400, 192 kHz, 24 bit), the generated pure tones were amplified (Rotel power amplifier, model RB-1050) and presented to the bats.

### Analysis of LFP data

The analysis was implemented using custom-written Matlab scripts (MATLAB R2015b, The Math Works Inc., MA, USA). All vocalizations were assessed offline using the Avisoft SAS Lab Pro software (v.5.2 Avisoft Bioacoustics, Germany). The initial acoustic trigger, non-echolocation calls (typical power maximum around 5-50 kHz) and echolocation calls (peaking above 50 kHz)^13^ were manually located, individually labelled, and their timing was exported to Matlab. To evade response contamination by other auditory stimuli, the “good” non-echolocation and echolocation calls were identified, which comprised at least 500 ms without any vocalization prior and following call production. The spectrograms of the vocalizations were calculated with a frame width of 0.8, a frame shift of 0.05 and a hamming window of 2048-points length.

To investigate LFPs during each task, the electrophysiological signal was filtered between 1-90 Hz (2nd order Butterworth filter), the line noise removed using the *rmlinesmovingwinc* function of the Chronux toolbox ^48^ and down-sampled from 20 kHz to 1 kHz. Additionally, the signals were normalized by calculating the z-score at each time point by subtracting the mean and dividing by the standard deviation per recording. Z-scoring was conducted across channels for the FAF (to respect amplitude relationships across channels) and for each channel of the CN individually.

To extract LFP fluctuations linked to vocalization production a randomization procedure was used. This randomization procedure rendered 10,000 non-echolocation and echolocation signals for the CN and each recording channel of the FAF. Each randomization trial was obtained by averaging 100 randomly chosen LFPs corresponding to either the non-echolocation or echolocation conditions. Note that because of extensive averaging, this randomization procedure removes signal components that are not locked to the vocalizations.

Time-frequency analysis was conducted for each randomization trial using the Chronux function *mtspecgramc* with 250 ms window size, 0.5 ms time step and a time-bandwidth product of 2, with 3 tapers. To compute the difference in power during the production of different call types, the logarithmic power spectrogram of the non-echolocation condition was subtracted from the logarithm of the power spectrogram obtained during echolocation. Statistical significance was evaluated using Cliff’s Delta (*d*), which was calculated based on the area under the receiver operating characteristic curve (AUROC, linear relation: Cliff’s delta = 2 * AUROC − 1) (Hentschke-“Measures of Effect Size”-toolbox). This measure ranges between −1 to 1 with almost identical observations rendering *d*-values around zero. The *d*-value borders for defining large, medium and small effect sizes were set to, 0.474, 0.333 and 0.147, respectively ^29^.

A binary support vector machine (SVM) classifier was used for predicting vocal output using the average spectral signal in each LFP band before vocalization production. The SVM classifier was trained (*fitcsvm* function, rbf kernel, Matlab 2015, single training, no standardization, fitting posterior probabilities after model creation) using signals obtained in 10,000 randomization trials (5,000 per vocalization type, see preceding text). SVM models obtained were cross-validated using 10-fold cross-validation. In a second step, labels were swapped in the training set before classification to assess the performance of the models in the absence of reliable training information. To assess statistical significance, the binominal distribution formula was calculated and the computed probability values of the data were compared to the chance of obtaining the same results by chance. The significance level was defined with p<0.05 which correspond to <5 %^49^.

To evaluate the oscillatory coherence and phase consistency between signals in the striatum and the different cortical depths of the FAF, the Chronux function *cohgramc* with the same parameters used for spectral analysis (see neural spectrogram specifications above) was used. This operation performed coherency calculations between all possible pairs of different channels in the FAF and each channel in the striatum (here, no randomization was used; in other words, we used the LFPs linked to the production of each echolocation and non-echolocation trial). Then, the average coherogram obtained between FAF channels and each striatal channel was calculated. For displaying and assessing the strength of coherency, the magnitude of the coherency (“coherence”) was used. Coherence values exceeding the 95th percentile of all coherence values obtained were labelled as significant.

Using the same pre-processing methods described above (filtering, down-sampling, z-scoring per recording and demeaning), LFP responses obtained from the frequency tuning paradigm were quantified. The absolute value of the analytical signal (obtained after Hilbert transforming) was used to calculate the instantaneous energy of each recording channel in response to each sound frequency tested. The frequency eliciting the highest amount of energy was labelled as best frequency.

### Analysis of spike data

Spiking activity was acquired by filtering neural signals in the frequency range of 300-3000 Hz (2nd order Butterworth filter). Spike detection was performed using the SpyKING CIRCUS toolbox with automatic clustering and a threshold of 5 median absolute deviations using the best spiking template per channel and recording ^50^. With a bin-size of 3ms, peri-stimulus time histograms (PSTHs) were calculated for both brain structures.

To investigate the relationship between spikes and LFPs, phase-locking values were calculated using the circular statistics toolbox ^51^. For phase locking calculations, only the time window before vocalization production was considered. The procedure used for calculating phase locking values is illustrated in Supplementary Figure 6 for one example echolocation trial. After extracting spike times and raw LFPs related to the isolated vocalization trial, the LFP signal was filtered in different frequency bands (e. g. theta (4-8 Hz), alpha (8-12 Hz), low beta (12-20 Hz), high beta (20-30 Hz), low gamma (30-50 Hz), and high gamma (50-80 Hz). Filtered LFPs were Hilbert-transformed and their instantaneous phase information was extracted. The phase at which spiking occurred for each LFP frequency band was stored and analysed using circular statistics (see below).

Circular distributions of LFP phases at which spiking occurred for each frequency band were compared with random-phase distributions obtained by extracting LFP phases at random time points not related to spiking. To obtain robust circular spike-phase and random-phase distributions, circular distributions were calculated 10,000 times, with 100 randomly chosen spike-phase and random-phase values included in each randomization trial. Two parameters were extracted from the circular distributions obtained in each spike- and random-phase trial: the distribution’s vector strength (VS, *circ_r* function in the circular statistics toolbox^51^) and its angular mean (*circ_mean* function in the circular statistics toolbox^51^). VS values obtained from all randomization trials were used for assessing statistical significance when comparing spike-phase and random-phase distributions using Bonferroni corrected Wilcoxon rank-sum tests (p<0.001). Angular mean values were used for visual display and for calculating population VS differences (dVS). In our calculations, positive dVS values indicate higher vector strength (VS) in the spike-phase distribution when compared to the random-phase control.

### Histological verification of striatal recordings

For visualization of the electrode implantation location, histological analysis was performed following the completion of the experiments. To locate the tracks of the chronically implanted tetrode in the striatum, an electric lesion was performed for 10 sec with 10 μA DC current using a Stimulus Isolator A365 (World Precision Instruments, USA) under deep anaesthesia prior to perfusion. Electric lesions were set for each animal on the last experimental day on the most ventral and dorsal striatal electrodes. Subsequently, the animals were euthanized with an intraperitoneal injection of 0.1 ml sodium pentobarbital (160 mg/ml, Narcoren, Boehringer-Ingelheim, Germany) and transcardially perfused using a peristaltic pump (Ismatec, Wertheim, Germany) with a pressure rate of 3-4 ml/min. The bats were perfused with 0.1 M phosphate buffer saline for 5 min, followed by a 4% paraformaldehyde solution for 30 min. After removing the surrounding tissue, muscles and skull, the brain was carefully eviscerated, fixed in 4% paraformaldehyde at 4°C for at least one night and placed in an ascending sucrose sequence solution (1 h in 10%, 2-3 h in 20%, 1 night in 30%) at 4°C to avoid the formation of ice crystals in the tissue. Subsequently, the brain was frozen in an egg yolk embedding encompassing the fixation in glutaraldehyde (25%) with CO_2_. For sectioning the frozen brain, a cryostat (Leica CM 3050S, Leica Microsystem, Wetzlar, Germany) was utilized and coronal slices (50μm thick) were prepared, mounted on gelatin-coated slides and Nissl stained. In brief, the brain slices were immersed in 96% ethanol overnight and 70% ethanol (5min), hydrated in distilled water (3×3min), stained in 0.5% cresylviolet (10min), rinsed in diluted glacial acetic acid (30sec), differentiated in 70% ethanol + glacial acetic acid until neuronal somata were still red-violet stained with only faint coloration of the background, fixed in an ascending alcohol sequence (2×5min in 96% ethanol, 2×5min in 100% isopropyl alcohol), cleaned by Rotihistol I, II and III solution (Carl-Roth GmbH, Karlsruhe, Germany) and covered with DPX mounting medium. The inspection of the lesion was facilitated by a bright-field, fluorescence microscope (Keyence BZ-9000, Neu-Isenburg, Germany). A Nissl staining of a bat brain with the associated track of a chronically implanted HQ4 tetrode in the dorsal part of the CN can be found in the in Supplementary Fig. 1d.

## Acknowledgments

This work was funded by the Deutsches Forschungsgemeinschaft (DFG) grant number HE7478/1-1. We are thankful you Gisa Prange for histological support, Christin Reißig for animal care and Manfred Kössl for valuable comments on this study.

## Author Contributions

JCH, FGR and KW designed the study. KW collected and analyzed the data. KW and JCH wrote the manuscript. KW, JCH and FGR revised the manuscript.

## References

1. Wegdell, F., Hammerschmidt, K. & Fischer, J. Conserved alarm calls but rapid auditory learning in monkey responses to novel flying objects. Nat. Ecol. Evol. 3, 1039–1042 (2019).

2. Kanwal, J. S. & Rauschecker, J. P. Auditory cortex of bats and primates: managing species-specific calls for social communication. Front. Biosci. 12, 4621–4640 (2007).

3. Patterson, R. D., Uppenkamp, S., Johnsrude, I. S. & Griffiths, T. D. The Processing of Temporal Pitch and Melody Information in Auditory Cortex. Neuron 36, 767–776 (2002).

4. Beetz, M. J., García-Rosales, F., Kössl, M. & Hechavarría, J. C. Robustness of cortical and subcortical processing in the presence of natural masking sounds. Sci. Rep. 8, 6863 (2018).

5. Park, H., Ince, R. A. A., Schyns, P. G., Thut, G. & Gross, J. Frontal Top-Down Signals Increase Coupling of Auditory Low-Frequency Oscillations to Continuous Speech in Human Listeners. Curr. Biol. 25, 1649–1653 (2015).

6. Grinnell, A. D. The neurophysiology of audition in bats: intensity and frequency parameters. J. Physiol. 167, 38–66 (1963).

7. Kothari, N. B., Wohlgemuth, M. J. & Moss, C. F. Dynamic representation of 3D auditory space in the midbrain of the free-flying echolocating bat. 1–29 (2018).

8. Kawasaki, M., Margoliash, D. & Suga, N. Delay-tuned combination-sensitive neurons in the auditory cortex of the vocalizing mustached bat. J. Neurophysiol. 59, 623–635 (1988).

9. Metzner, W. A possible neuronal basis for Doppler-shift compensation in echo-locating horseshoe bats. Nature 341, 529–532 (1989).

10. Giraud, A.-L. et al. Endogenous Cortical Rhythms Determine Cerebral Specialization for Speech Perception and Production. Neuron 56, 1127–1134 (2007).

11. Smith, A. & Denny, M. High-frequency oscillations as indicators of neural control mechanisms in human respiration, mastication, and speech. J. Neurophysiol. 63, 745–758 (1990).

12. Teeling, E. C. et al. Molecular evidence regarding the origin of echolocation and flight in bats. Nature 403, 188–192 (2000).

13. Hechavarría, J. C., Beetz, M. J., Macias, S. & Kössl, M. Distress vocalization sequences broadcasted by bats carry redundant information. J. Comp. Physiol. A 202, 503–515 (2016).

14. Voorn, P., Vanderschuren, L. J. M. J., Groenewegen, H. J., Robbins, T. W. & Pennartz, C. M. A. Putting a spin on the dorsal–ventral divide of the striatum. Trends Neurosci. 27, 468–474 (2004).

15. Birba, A. et al. Losing ground: Frontostriatal atrophy disrupts language embodiment in Parkinson’s and Huntington’s disease. Neurosci. Biobehav. Rev. 80, 673–687 (2017).

16. Leh, S. E., Ptito, A., Chakravarty, M. M. & Strafella, A. P. Fronto-striatal connections in the human brain: A probabilistic diffusion tractography study. Neurosci. Lett. 419, 113–118 (2007).

17. Ferry, A. T., Öngür, D., An, X. & Price, J. L. Prefrontal cortical projections to the striatum in macaque monkeys: Evidence for an organization related to prefrontal networks. J. Comp. Neurol. 425, 447–470 (2000).

18. Radulescu, E. et al. Abnormalities in fronto-striatal connectivity within language networks relate to differences in grey-matter heterogeneity in Asperger syndrome. NeuroImage Clin. 2, 716–726 (2013).

19. Vargha-Khadem, F., Gadian, D. G., Copp, A. & Mishkin, M. FOXP2 and the neuroanatomy of speech and language. Nat. Rev. Neurosci. 6, 131 (2005).

20. Robles, S. G., Gatignol, P., Capelle, L., Mitchell, M.-C. & Duffau, H. The role of dominant striatum in language: a study using intraoperative electrical stimulations. J. Neurol. Neurosurg. & Psychiatry 76, 940 LP–946 (2005).

21. Holland, R. et al. Speech Facilitation by Left Inferior Frontal Cortex Stimulation. Curr. Biol. 21, 1403–1407 (2011).

22. Schwartz, C. P. & Smotherman, M. S. Mapping vocalization-related immediate early gene expression in echolocating bats. Behav. Brain Res. 224, 358–368 (2011).

23. Eiermann, A. & H Esser, K. Auditory responses from the frontal cortex in the short-tailed fruit bat Carollia perspicillata. Neuroreport 11, (2000).

24. Kanwal, J. S., Gordon, C. A. M., Peng, J. P. & Heinz-esser, K. Auditory responses from the frontal cortex in the mustached bat, Pteronotus parnellii. 11, 367–372 (2000).

25. Kobler, J. B., Isbey, S. F. & Casseday, J. H. Auditory pathways to the frontal cortex of the mustache bat, Pteronotus parnellii. Science (80-.). 236, 824 LP–826 (1987).

26. Laubach, M., Amarante, L. M., Swanson, K. & White, S. R. What, If Anything, Is Rodent Prefrontal Cortex? eneuro 5, ENEURO.0315-18.2018 (2018).

27. Seamans, J. K., Lapish, C. C. & Durstewitz, D. Comparing the prefrontal cortex of rats and primates: Insights from electrophysiology. Neurotox. Res. 14, 249–262 (2008).

28. Bastos, A. M., Loonis, R., Kornblith, S., Lundqvist, M. & Miller, E. K. Laminar recordings in frontal cortex suggest distinct layers for maintenance and control of working memory. Proc. Natl. Acad. Sci. 115, 1117 LP–1122 (2018).

29. Romano, J., Kromrey, J. D., Coraggio, J. & Skowronek, J. Appropriate statistics for ordinal level data: Should we really be using t-test and Cohen’sd for evaluating group differences on the NSSE and other surveys. in annual meeting of the Florida Association of Institutional Research 1–33 (2006).

30. Kramer, M. A. An Introduction to Field Analysis Techniques: The Power Spectrum and Coherence. (2013).

31. Zhang, W. & Yartsev, M. M. Correlated Neural Activity across the Brains of Socially Interacting Bats. Cell (2019). doi:10.1016/j.cell.2019.05.023

32. Yartsev, M. M., Witter, M. P. & Ulanovsky, N. Grid cells without theta oscillations in the entorhinal cortex of bats. Nature 479, 103 (2011).

33. Eliav, T. et al. Nonoscillatory Phase Coding and Synchronization in the Bat Hippocampal Formation. Cell 175, 1119–1130.e15 (2018).

34. Arnal, L. H., Doelling, K. B. & Poeppel, D. Delta–Beta Coupled Oscillations Underlie Temporal Prediction Accuracy. Cereb. Cortex 25, 3077–3085 (2014).

35. Wiener, M., Parikh, A., Krakow, A. & Coslett, H. B. An Intrinsic Role of Beta Oscillations in Memory for Time Estimation. Sci. Rep. 8, 7992 (2018).

36. Engel, A. K. & Fries, P. Beta-band oscillations—signalling the status quo? Curr. Opin. Neurobiol. 20, 156–165 (2010).

37. Khanna, P. & Carmena, J. M. Beta band oscillations in motor cortex reflect neural population signals that delay movement onset. Elife 6, e24573 (2017).

38. Bartolo, R., Prado, L. & Merchant, H. Information Processing in the Primate Basal Ganglia during Sensory-Guided and Internally Driven Rhythmic Tapping. J. Neurosci. 34, 3910 LP–3923 (2014).

39. Schroeder, C. E. & Lakatos, P. Low-frequency neuronal oscillations as instruments of sensory selection. Trends Neurosci. 32, 9–18 (2009).

40. von Stein, A. & Sarnthein, J. Different frequencies for different scales of cortical integration: from local gamma to long range alpha/theta synchronization. Int. J. Psychophysiol. 38, 301–313 (2000).

41. Buzsaki, G, D. A. Neuronal oscillations in cortical networks. Science (80-.). 304, 1926–9 (2004).

42. Lepski, G., Arévalo, A., Valle, A. C., Ballester, G. & Gharabaghi, A. do Increased coherence among striatal regions in the theta range during attentive wakefulness. Brazilian J. Med. Biol. Res. = Rev. Bras. Pesqui. medicas e Biol. 45, 763–770 (2012).

43. Harris, K. D. & Mrsic-flogel, T. D. Cortical connectivity and sensory coding. doi:10.1038/nature12654

44. Lewandowski, B. C. & Schmidt, M. Short Bouts of Vocalization Induce Long-Lasting Fast Gamma Oscillations in a Sensorimotor Nucleus. J. Neurosci. 31, 13936 LP–13948 (2011).

45. Fries, P. Rhythms for Cognition: Communication through Coherence. Neuron 88, 220–235 (2015).

46. Medvedev, A. V & Kanwal, J. S. Communication call-evoked gamma-band activity in the auditory cortex of awake bats is modified by complex acoustic features. Brain Res. 1188, 76–86 (2008).

47. van der Meer, M. A. A. & Redish, A. D. Covert Expectation-of-Reward in Rat Ventral Striatum at Decision Points. Front. Integr. Neurosci. 3, 1 (2009).

48. Andrews, P., Loader, C. & Shukla, R. The Chronux Manual. (2008).

49. Quian Quiroga, R., Kraskov, A., Mormann, F., Fried, I. & Koch, C. Single-Cell Responses to Face Adaptation in the Human Medial Temporal Lobe. Neuron 84, 363–369 (2014).

50. Yger, P. et al. A spike sorting toolbox for up to thousands of electrodes validated with ground truth recordings in vitro and in vivo. Elife 7, e34518 (2018).

51. Berens, P. CircStat: A MATLAB Toolbox for Circular Statistics. J. Stat. Software; Vol 1, Issue 10 (2009). doi:10.18637/jss.v031.i10

