## supplementary figures S1-S9 for "Fronto-striatal oscillations predict vocal output in bats"

**Supplementary Information**

**Supplementary Figures 1-9**

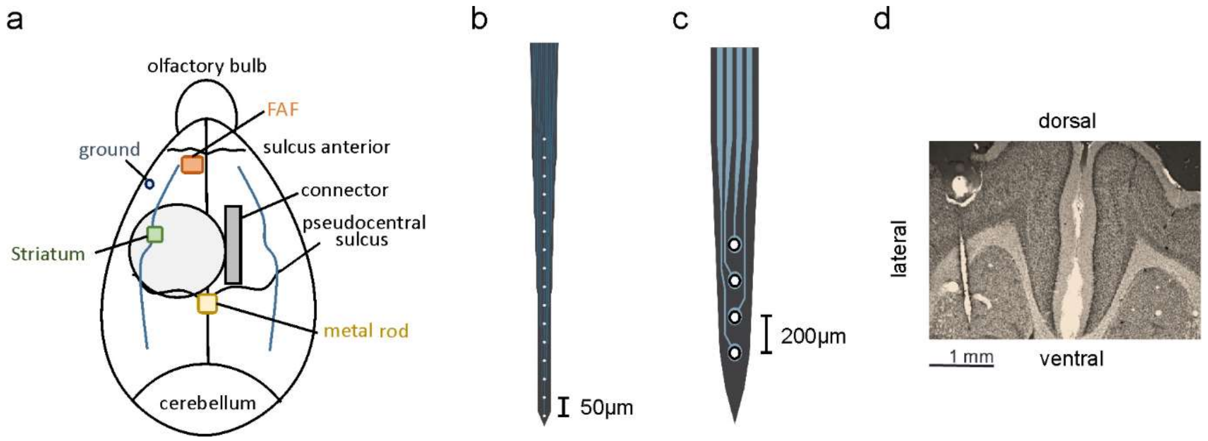


**Supplementary Figure S1 Electrode implantation procedure. a,** Schematic outline of the implantation sites. Olfactory bulb denotes the anterior part of the brain while the cerebellum is found in the posterior part. FAF= frontal auditory field. **b,** Mapping of the A16 laminar silicon probe with 50 µm spacing between electrodes (which was implanted in the FAF) and **c,** the HQ4 laminar tetrode with 200 µm between recording sites chronically implanted in the CN. **d**, Nissl stained section including the track of an HQ4 laminar tetrode implanted in the caudate nucleus (4x magnification).

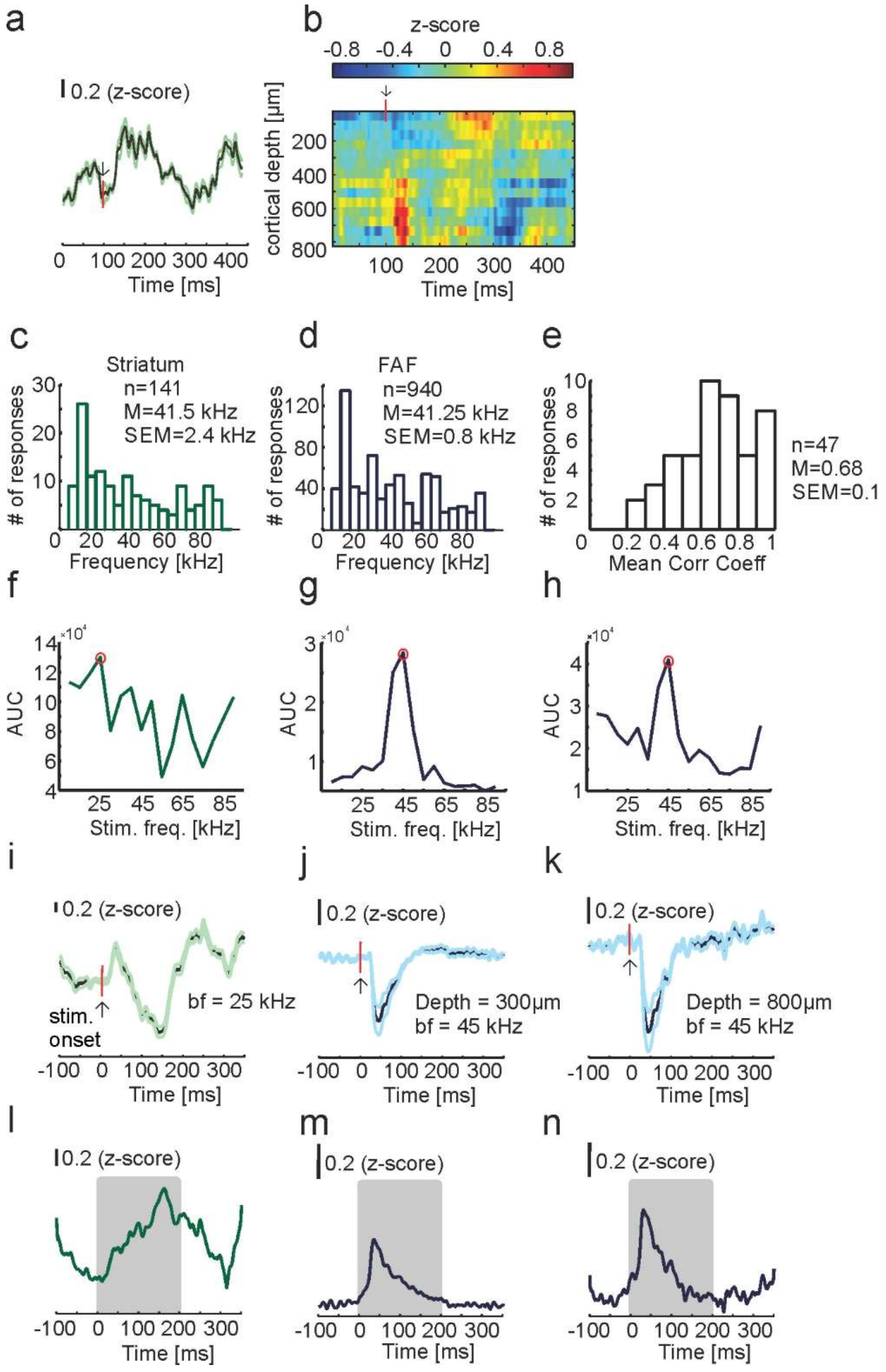


**Supplementary Figure S2. CN and FAF display auditory responsiveness to stimulation with pure tones. a**, Population mean striatal activity ± SEM in response to the best frequency (bf) across recording sites. The arrow indicates stimulus onset. **b**, Bitmap of the amplitude of z-scored LFPs from 100 ms before up to 350 ms after the stimulus onset across cortical depths in the FAF. **c,** Histogram of bfs in the striatum. Mean (M) and standard error of the mean (SEM) are indicated. **d**, Histogram of bfs in the FAF. Both brain structures exhibited pronounced auditory responsiveness to low frequencies. **e,** Distribution of the mean correlation coefficient obtained by correlating tuning curves of all simultaneous recordings in different FAF depths. The mean value across recordings (n=47) was calculated. The high mean correlation (0.68) indicates similar tuning in neighboring channels. **f,** Exemplary area under the curve (AUC) of the LFP response to different simulation frequencies in the striatum and the FAF at depths of 300 µm (**g**), and 800 µm (**h**). The red circles indicate the bf. **i, j, k** Mean LFP traces (±SEM) following auditory stimulation at the bf in the same exemplary recordings mentioned above. The arrows refer to the stimulation onset. Subpanels **l, m, n** show the instantaneous energy of the respective LFP traces shown in i-k.

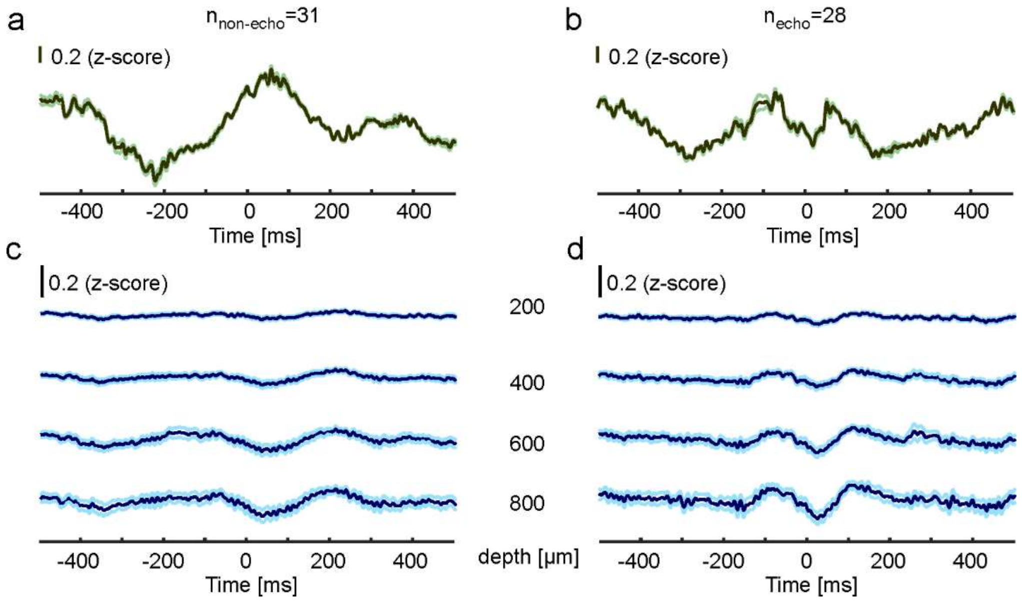


**Supplementary Figure S3. Differences in LFP activity during vocalization production in the FAF and CN in an illustrative paired recording. a,** Mean LFP (±SEM) of all isolated non-echolocation calls (n=31) during one recording in the striatum and **b,** of all isolated echolocation calls in the same recording (n=28). **c**, The mean ± SEM traces in the FAF at four representative depths (200, 400, 600 and 800 µm) in the same time period as **a** for non-echolocation and **d**, echolocation. In the FAF, highest differences between call types occurred in the deepest channels. The example traces show activity before call onset, which could be used to predict the type of vocal output in both brain regions.

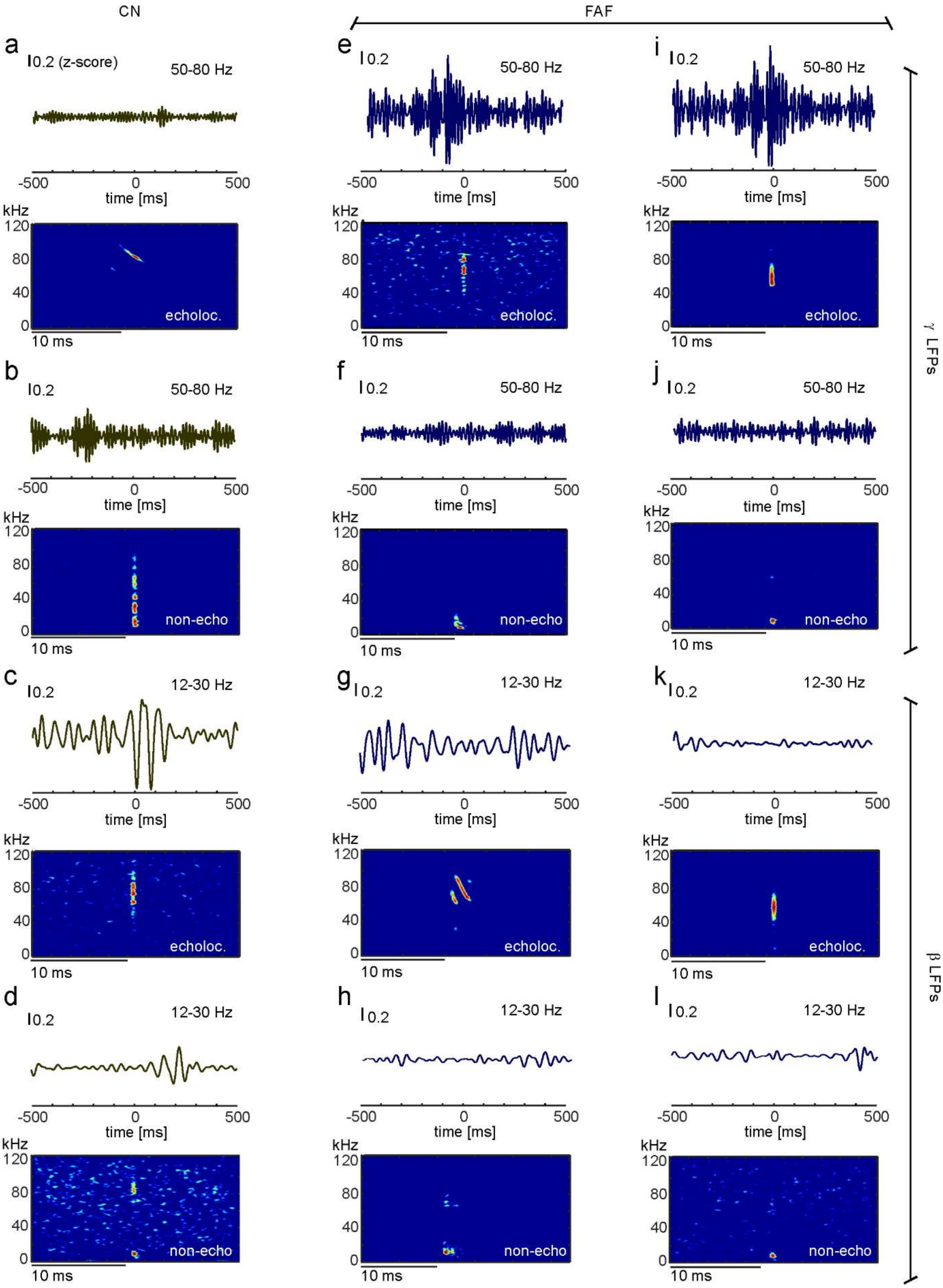
CN
FAF
a
0.2 (z-score)
50-80 Hz
-500
0
500
time [ms]
kHz
120
80
40
0
echoloc.
10 ms
b
0.2
50-80 Hz
non-echo
c
0.2
12-30 Hz
d
0.2
12-30 Hz
non-echo
e
f
g
h
i
j
k
l
γ LFPs
β LFPs

**Supplementary Figure S4. Example neural recordings showing different LFP frequency patterns in the FAF and CN.** In each subpanel, bottom panels show the broadcasted call, whereas top panels show filtered LFP traces obtained before and after call production in each case. **a**, Individual example LFP in the CN during echolocation and **b**, non-echolocation filtered in high gamma (50-80 Hz). **c**, Example filtered LFPs (in the beta range) during echolocation and **d**, non-echolocation. **e - h,** Analogous exemplification for the FAF at 800 µm cortical depth. **i - l**, A second example FAF recording during echolocation (i and k) and non-echolocation trials (j and l). In the FAF, during echolocation trials, high gamma and beta power occurs before call production.

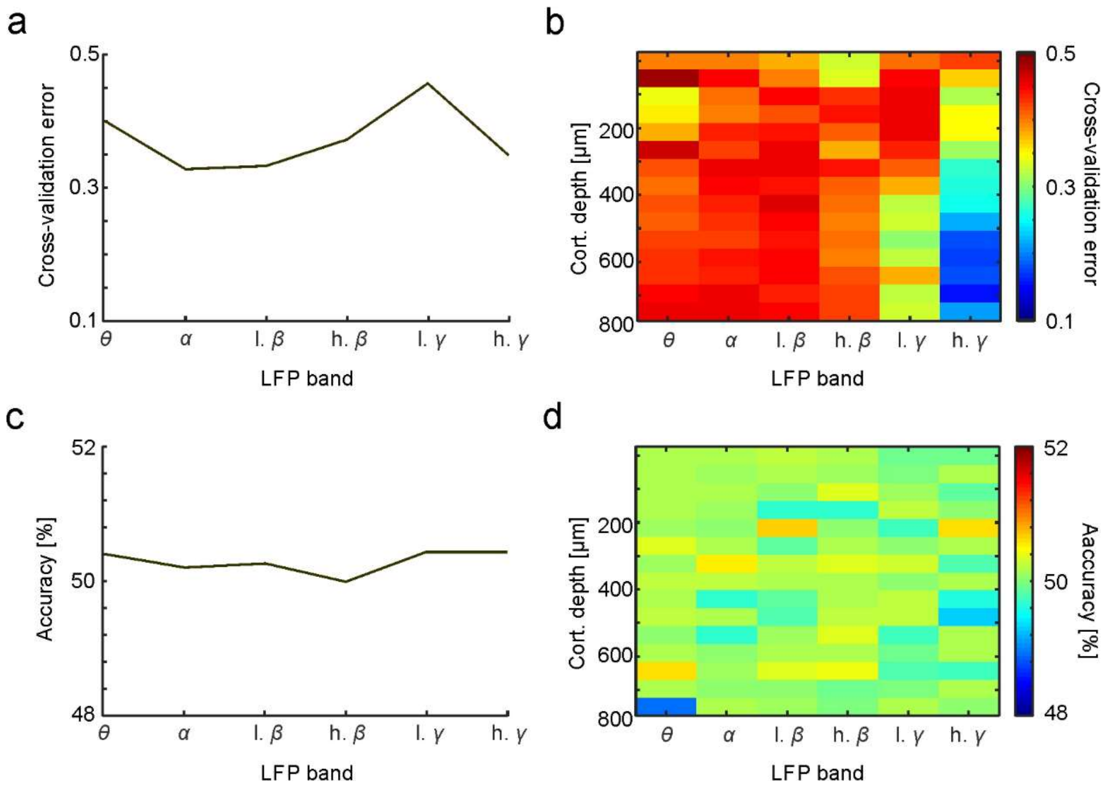


**Supplementary Figure S5. Cross-validation and prediction accuracy of support vector machine prediction models. a**, Cross-validation error across LFP frequencies in the CN and **b**, in the FAF. Note that the lowest cross-validation errors occurred in the deep channels of the FAF in high gamma. **c,** Assessment of the SVM model performance based on randomly chosen labels in the training sessions in the CN and **d**, in the FAF. With insufficient training information, model accuracy drops to values around chance level ~50%.

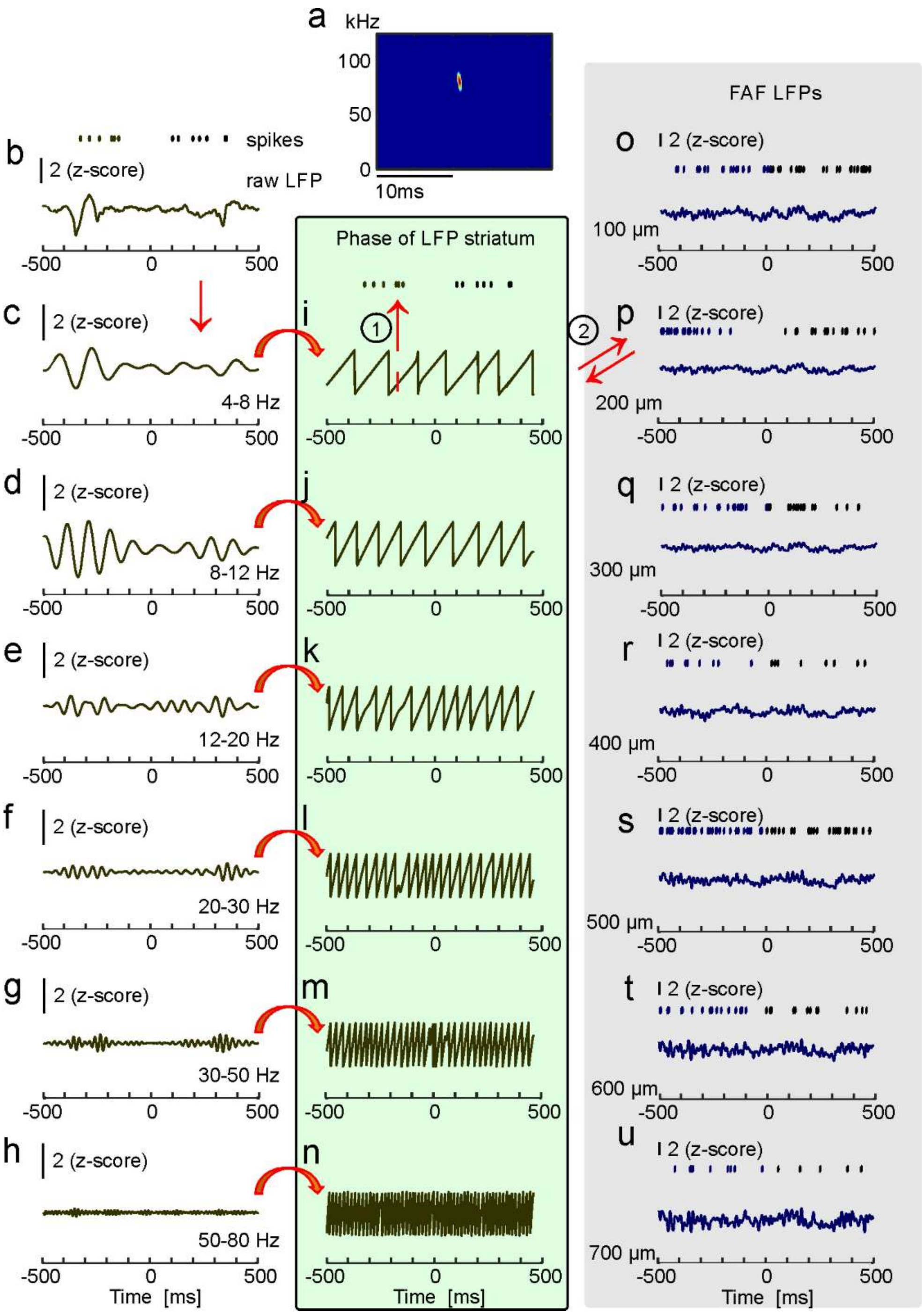


**Supplementary Figure S6. Illustrative calculation of phase locking values in one example trial within and between brain structures. a,** Spectrogram of the echolocation call emitted in this vocalization trial. **b,** Top: spike times obtained in this trial in the CN (represented as dots). Bottom: simultaneously recorded raw LFP trace in the CN. Filtered

LFP signals are shown in **c**: theta, **d:** alpha, **e:** low-beta, **f:** high-beta, **g:** low-gamma and **h:** high-gamma. **i-n,** Instantaneous phase values extracted after Hilbert transforming the filtered LFP signals associated to panel c-h, respectively. Phase values at the time points in which spiking occurred were used for phase-locking calculations. **o-u** Recorded spike times (top, blue: spikes before call onset) in the FAF were correlated with the phase of the LFP within the same cortical depth (here: raw traces at 100-700 µm, in steps of 100 µm, are shown) and to the LFP phase obtained simultaneously in the CN, and vice versa. This procedure rendered inter-area and intra-area phase locking values. The computed values were used as a measure of how strong the spike events were linked to the phase of LFPs within one brain region and in the fronto-striatal network.

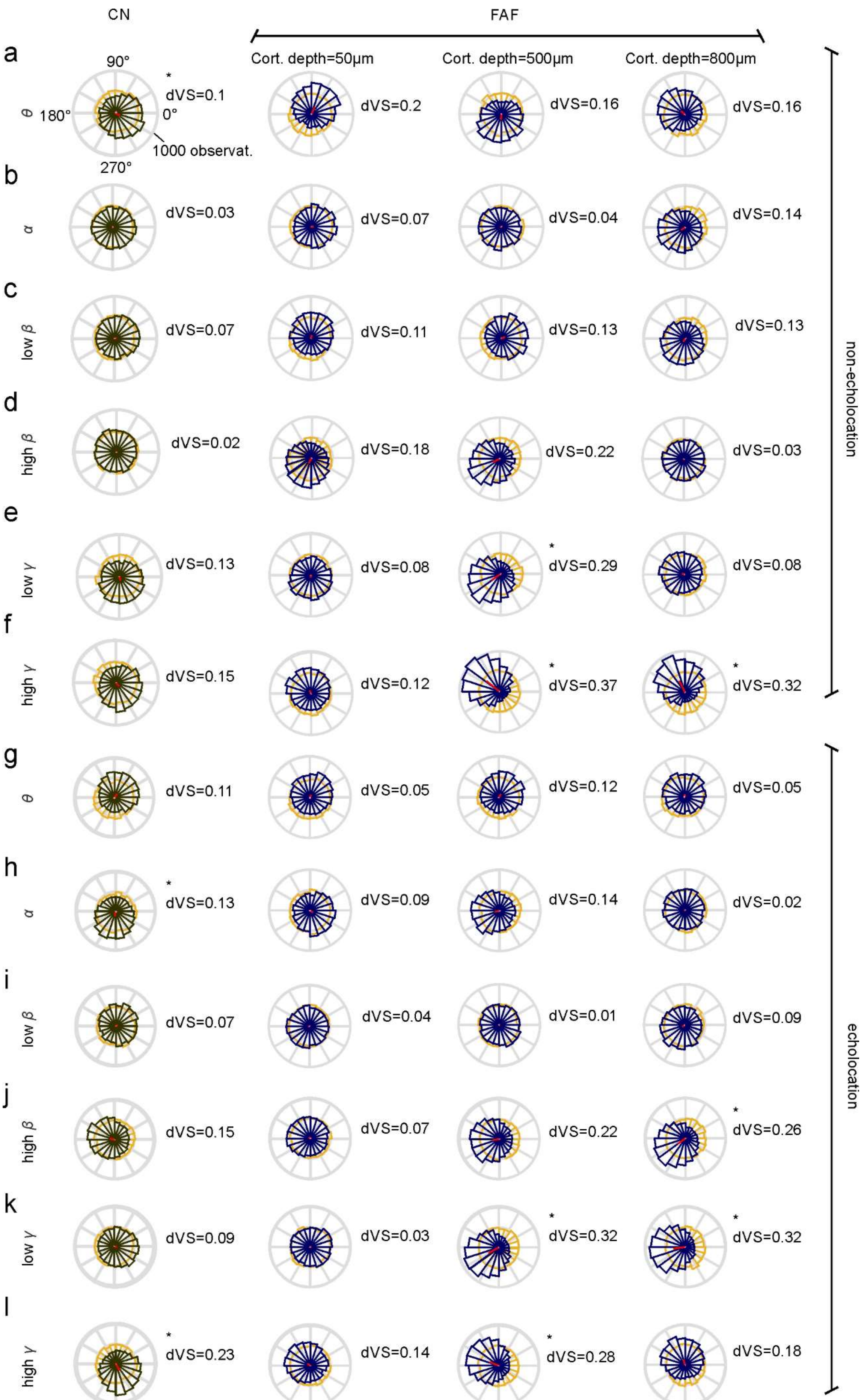

CN
FAF
Cort. depth=50µm
Cort. depth=500µm
Cort. depth=800µm
a
θ
90°
180°
0°
270°
1000 observat.
* dVS=0.1
dVS=0.2
dVS=0.16
dVS=0.16
b
α
dVS=0.03
dVS=0.07
dVS=0.04
dVS=0.14
c
low β
dVS=0.07
dVS=0.11
dVS=0.13
dVS=0.13
d
high β
dVS=0.02
dVS=0.18
dVS=0.22
dVS=0.03
e
low γ
dVS=0.13
dVS=0.08
* dVS=0.29
dVS=0.08
f
high γ
dVS=0.15
dVS=0.12
* dVS=0.37
* dVS=0.32
non-echolocation
g
θ
dVS=0.11
dVS=0.05
dVS=0.12
dVS=0.05
h
α
* dVS=0.13
dVS=0.09
dVS=0.14
dVS=0.02
i
low β
dVS=0.07
dVS=0.04
dVS=0.01
dVS=0.09
j
high β
dVS=0.15
dVS=0.07
dVS=0.22
* dVS=0.26
k
low γ
dVS=0.09
dVS=0.03
* dVS=0.32
* dVS=0.32
l
high γ
* dVS=0.23
dVS=0.14
* dVS=0.28
dVS=0.18
echolocation

**Supplementary Figure S7. Circular mean distributions illustrating spike-phase locking in the CN and in example FAF channels in different LFP frequency bands. a-f,** Circular mean distributions for non-echolocation in theta (**a**) alpha (**b**), low beta (**c**), high beta (**d**), low gamma (**e**) and high gamma (**f**). In each column**,** the first row indicates values acquired from the CN, whereas columns 2-4 display data obtained at three different FAF depths (i.e. 50 µm, 500 µm and 800 µm). Red lines indicate the vector strength (VS) of each circular distribution. Data plotted in orange represent surrogate distributions. **g- l** Same as a-h but for the echolocation condition. dVS = difference in vector strength between the spike-phase locking distributions and the surrogate distributions; * $p<0.001$ (Wilcoxon rank-sum comparing VS values across randomization trials, Bonferroni corrected).

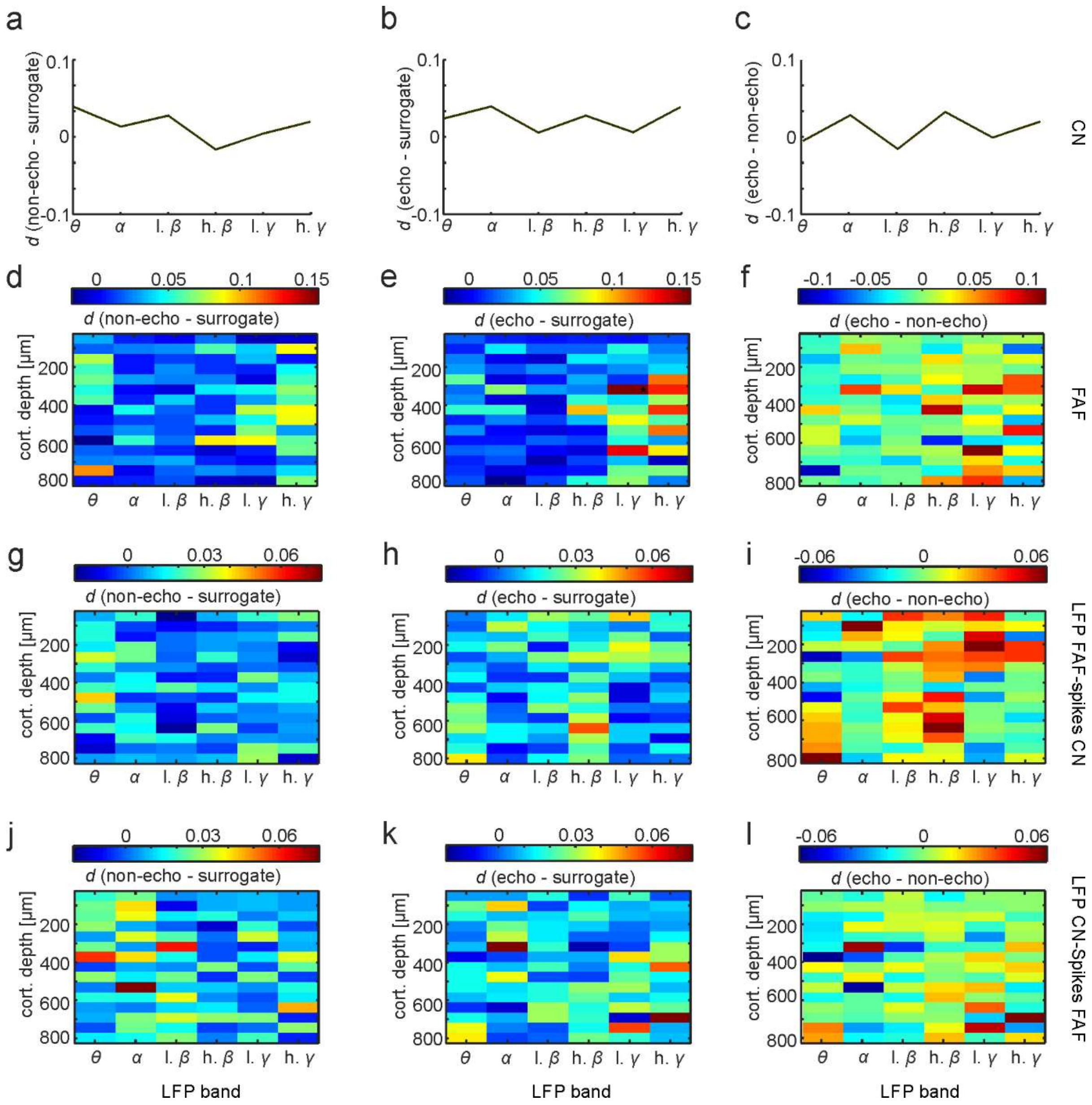


**Supplementary Figure S8. Effect sizes of comparisons in phase locking between the types of vocal output within and between fronto-striatal areas obtained using Cliff's Delta (*d*). a,** *d*-values obtained when comparing vector strengths (VS) in the non-echolocation condition to the surrogate distributions in the CN across LFP frequencies bands (n=10,000; l. =low; h.=high)**. b,** Same as **a**, for the echolocation - surrogate condition. **c,** *d*-values obtained when comparing the VS of the echolocation and non-echolocation conditions (positive = higher phase locking before echolocation; negative = higher phase locking before non-echolocation). **d-f,** *d*-values across depths and LFP frequency bands in the FAF for the non-echolocation – surrogate condition, **e**, echolocation – surrogate and **f**, echolocation - non-echolocation. **g-i** Equivalent comparisons to the ones in d-f when linking the phase of LFPs in the FAF to the simultaneously recorded spikes in the CN. **j-l** Same as g-i for the reversed situation (spikes of the FAF linked to the phase of LFPs in the CN). * indicate a small effect size (i.e. $d$>0.147, found only in one case, see panel e).

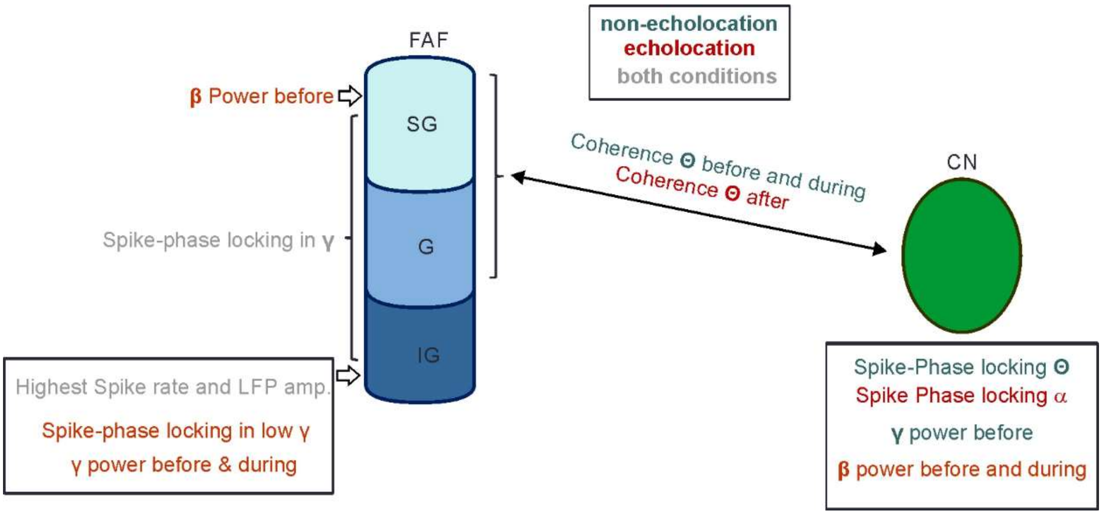


**Supplementary Figure S9. Visual abstract depicting the main results presented in this study.** FAF: frontal auditory field, CN: caudate nucleus, SG, G, and IG: indicate a putative subdivision of the FAF into supragranular, granular and infragranular layers, respectively. Note that we do not report data on the directionality of the connection between both regions and thus, functional coupling is displayed with a double arrow. Different electrophysiological parameters such as LFP power measurements, spike-phase locking and LFP-LFP coherence demonstrate that fronto-striatal circuits can predict ensuing vocal output in bats.
